# LSm4 biomolecular condensates drive XRN2-mediated RNA decay at DNA double-strand breaks to facilitate repair

**DOI:** 10.64898/2026.08.19.745694

**Authors:** Malak M. Darawshe, Laila A. Bishara, Enas R. Abu-Zhayia, Alma Sophia Barisaac, Feras E. Machour, Christeen Elmor, Nabieh Ayoub

## Abstract

Maintenance of genome integrity requires accurate repair of DNA double-strand breaks (DSBs), particularly within transcriptionally active regions. Persistent R-loops at DSBs can impede homologous recombination (HR) repair. While factors that resolve R-loops at DSB sites are known, the mechanisms ensuring timely degradation of nascent RNA to prevent pathological R-loop accumulation remain elusive. Here, we identified a critical role for the RNA-binding protein LSm4 in orchestrating localized RNA decay at DSBs to facilitate repair. We demonstrated that among LSm1-8 subunits, only LSm4 undergoes liquid-liquid phase separation (LLPS) and forms biomolecular condensates (BCs) specifically at DSBs in transcriptionally active chromatin. These damage-induced LSm4 BCs function as hubs that promote nuclear RNA decapping and recruit the 5′→3′ exonuclease XRN2 to degrade nascent transcripts proximal to DSBs. Accordingly, LSm4-XRN2 axis suppresses R-loop hyperaccumulation, thereby enabling efficient RAD51 filament assembly and intact HR repair. Consequently, loss of LSm4 increases translocations and leads to genomic instability. Collectively, our findings define a new regulatory layer in which LSm4 BCs spatially license RNA degradation, preventing R-loop accumulation at DSB microenvironment to facilitate error-free repair.

**Highlights:**

- LSm4 forms biomolecular condensates (BCs) via liquid-liquid phase separation (LLPS) at DSBs within transcriptionally active chromatin.
- LSm4 BCs create a hub for efficient RNA decay at DSB sites.
- LSm4 BCs promote nuclear RNA decapping and target XRN2 to DSBs to drive RNA decay.
- LSm4 suppresses R-loop accumulation to ensure intact homologous recombination repair of DSBs and maintain genomic stability.

## Introduction

Faithful repair of DNA double-strand breaks (DSBs) is a cornerstone of genomic stability. In eukaryotic cells, homologous recombination (HR) is a high-fidelity repair mechanism functioning during the S and G2 phases, that uses an undamaged sister chromatid as a template to restore the original DNA sequence ^1-4^. HR is particularly critical for accurate repair of DSBs occurring within transcriptionally active regions, where the preservation of coding sequence is vital to cellular fitness ^5–14^. Yet, active transcription machinery poses a formidable challenge to HR by physically impeding the access of repair factors to DSBs, as transcription complexes and nascent RNAs create a steric barrier that must be resolved ^15–19^. To mitigate this conflict, DSB induction triggers transient transcriptional silencing by modifying the histone code and inhibiting RNA Polymerase II (RNAPII) activity at DSB sites ^20–26^. However, pausing of RNAPII promotes re-annealing of nascent transcripts to the template DNA, leading to the formation of three-stranded DNA:RNA hybrid structures known as R-loops ^27–34^. Although R-loops fulfill certain regulatory roles, their persistence at DSBs directly impedes DNA end resection and interferes with the assembly of RAD51 recombinase filaments, thereby compromising HR efficiency and promoting genomic instability^30,33,35–41^. While cells utilize dedicated helicases like Senataxin (SETX) to resolve R-loops ^42,43^, the subsequent fate of the displaced and the nascent RNA transcripts at DSB sites is poorly understood. Timely degradation of nascent RNA nearby DSB sites may represent a fundamental step to prevent R-loop reformation and ensure efficient repair, yet the mechanisms responsible for clearing nascent RNA from DSB sites remain largely unexplored.

The LSm protein family comprises multiple RNA-binding proteins that share a conserved Sm motif ^44–46^. Eight of these LSm proteins are evolutionarily conserved across eukaryotes and assemble into two main complexes with distinct functions. The cytoplasmic LSm1-7 complex initiates mRNA decay by binding the 3ʹ ends of deadenylated transcripts and, together with Pat1, bridging the 3ʹ end to the 5ʹ cap to facilitate decapping by Dcp1/Dcp2. The decapped mRNA is then rapidly degraded by the 5ʹ→3ʹ exonuclease Xrn1 ^47–51^. In contrast, the nuclear LSm2-8 complex plays a central role in pre-mRNA splicing by stabilizing U6 snRNA and promoting assembly of U4/U6 and U4/U6·U5 snRNPs during spliceosome biogenesis^52,53^. One shared component of both complexes is the LSm4 subunit, which comprises 139 amino acids. Interestingly, LSm4 facilitates the assembly of processing bodies (P-bodies), cytoplasmic biomolecular condensates involved in mRNA storage, decay, and translational repression ^54^. Intriguingly, network analysis of the DNA repair interactome revealed LSm4 in proximity to several core DSB repair factors. However, the functional significance of this interaction remains entirely unexplored ^55^.

Here, we show that the RNA-binding protein LSm4 plays a previously unrecognized role in orchestrating RNA clearance at DNA DSBs through biomolecular condensate formation. We find that among the LSm1-8 family members, only LSm4 undergoes LLPS and is specifically recruited to DSBs occurring within transcriptionally active chromatin regions. At these damage sites, LSm4 assembles into dynamic condensates that promote nuclear RNA decapping and facilitate the recruitment of the 5′→3′ exonuclease XRN2, thereby driving efficient degradation of nascent transcripts proximal to DNA break sites. We demonstrate that this LSm4-XRN2 axis regulates R-loops homeostasis at DSBs and ensures intact HR repair. Collectively, our work establishes damage-induced LSm4 condensates as essential regulators of the DSB repair process and identifies RNA decay as a critical determinant of genome stability in transcriptionally active regions.

## Results

### Among LSm1-8 proteins, only LSm4 forms nuclear biomolecular condensates

To define the subcellular distribution of LSm4, we generated a U2OS cell line in which a Flag tag was endogenously knocked in at the C-terminus of the *LSm4* gene (Supplementary Figures 1A, 1B). Immunofluorescence analysis using an anti-Flag antibody revealed that LSm4 exhibits both cytoplasmic and nuclear localization, with notable enrichment in nucleolus. LSm4 nucleolar localization was confirmed by strong colocalization with the nucleolar marker Nucleolin (Figure 1A). Consistent with these observations, U2OS cells expressing an EGFP- LSm4 fusion protein exhibits a similar distribution pattern, suggesting that EGFP-LSm4 fusion faithfully recapitulates the localization of endogenous LSm4 (Figure 1B). Strikingly, among the human LSm1-8 proteins, only LSm4 exhibits nucleolar localization while the other LSm1- 8 proteins are excluded from the nucleolus (Figures 1B and Supplementary Figure 1C). Altogether, our data demonstrate that nucleolar condensation is a specific property of LSm4 that distinguishes it from other LSm1-8 proteins. Since nucleolar assembly is driven by liquid-liquid phase separation (LLPS) ^56^ and recombinant LSm4 protein undergoes LLPS *in vitro* ^57^, we sought to determine whether LSm4 forms condensates via LLPS in cells. First, to assess the liquid-like properties of LSm4 nucleolar condensates, we performed fluorescence recovery after photobleaching (FRAP) on EGFP-LSm4 at the nucleolus. Quantitative analysis revealed rapid fluorescence recovery, with a half-time (t₁/₂) of approximately one second, consistent with highly dynamic, liquid-like behavior of LSm4 (Figure 1C). Second, we employed an OptoDroplet assay using the blue light responsive photosensor Cry2WT to test the capacity of LSm4 to form condensates upon blue light activation ^58^. U2OS cells expressing either mCherry-labeled Cry2-pHR (Cry2WT) or LSm4-Cry2WT fusion were exposed to blue light and assessed for OptoDroplet formation. Strikingly, LSm4-Cry2WT forms rapid and pronounced OptoDroplet clusters both at the cytoplasm and the nucleus, whereas Cry2WT alone did not form clusters (Figure 1D, Supplementary Movies 1, 2). Moreover, neither LSm1 nor LSm8 formed OptoDroplet foci, consistent with their inability to form nucleolar condensates (Figure 1D, Supplementary Movies 3, 4). Together, these data provide the first evidence that LSm4, unlike other LSm subunits, undergoes LLPS and forms biomolecular condensates (BCs) in living cells.

**Figure 1:**
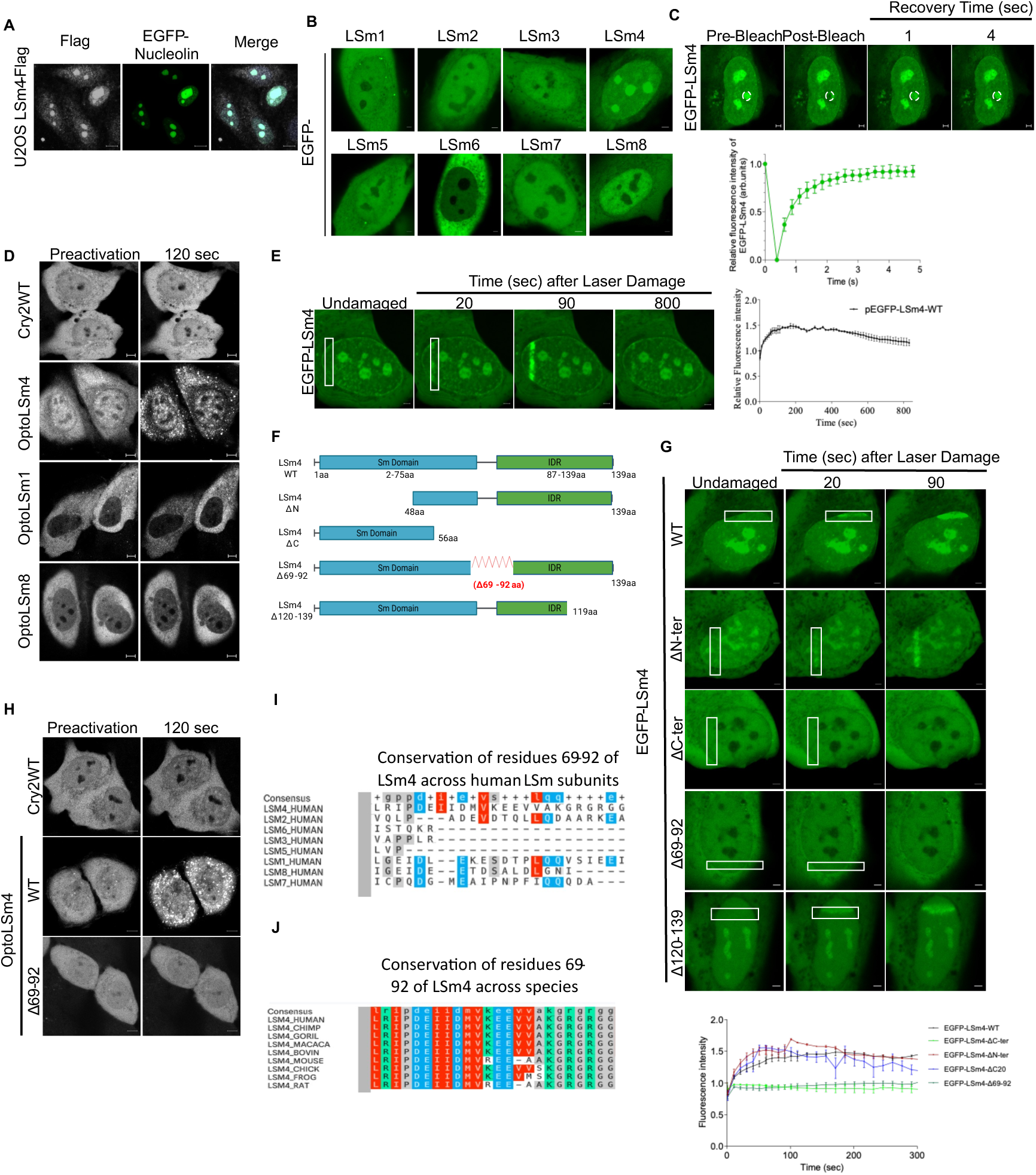
LSm4 forms nuclear biomolecular condensates at DNA damage sites via residues 69-92. (A) Representative immunofluorescence image showing LSm4 subcellular localization in U2OS^LSm4-Flag^ cells. Flag staining represents LSm4-Flag knock-in, and EGFP represents EGFP-Nucleolin. Scale bar 10 µm. (B) Representative confocal images showing the localization of EGFP-LSm1-8 proteins in U2OS cells. Scale bar 2 µm. (C) Top, representative images of U2OS cell expressing EGFP-LSm4 subjected to FRAP followed by time-lapse imaging post photobleaching. Scale bar 2 µm. Bottom, quantification of EGFP-LSm4 FRAP. Each data point represents the mean of 12 cells, and data are presented as mean ± standard deviation (SD). (D) Representative fluorescence images of U2OS cells expressing Cry2WT, OptoLSm4, OptoLSm1, and OptoLSm8, before and 120 seconds after light activation with 488nm laser. Scale bar 5 μm. (E) Left, Representative timelapse image of a U2OS cell expressing EGFP-LSm4 fusion and its localization at the indicated time points post laser- microirradiation (marked with white rectangle). Scale bar 2 µm. Right, quantification of EGFP- LSm4 relative fluorescence at laser-microirradiated sites. Data are presented as mean ±SD; n=4. (F) Schematic diagram showing the domain organization of full-length LSm4 (WT) and the corresponding deletion mutants. IDR: Intrinsically Disordered Region. (G) Top, timelapse images of the localization of EGFP-LSm4 WT and deletion mutants at laser-microirradiated sites (marked with white rectangle) in U2OS cells. Scale bar 2 µm. Bottom, quantification of fluorescence recovery after photobleaching for EGFP-LSm4 WT and deletion mutants. Data are presented as mean ±SD; n=3. (H) Representative fluorescence images of U2OS cells expressing Cry2WT, OptoLSm4^WT^ and OptoLSm4^Δ69–92^, before and 120 seconds after light activation with 488nm laser. Scale bar 5 μm. (I) Multiple sequence alignment of residues 69- 92 of human LSm4 with other human LSm proteins. (J) Multiple sequence alignment of residues 69-92 of human LSm4 across species.

### LSm4 forms biomolecular condensates at DNA damage sites via residues 69-92

Accumulating evidence indicates that several RNA-binding proteins, such as FUS, condensates at DNA damage sites in a PARP1-dependent manner ^59–64^. Given that LSm4 is also an RNA- binding protein, we asked whether it similarly undergoes condensation at DNA damage sites. We first monitored the localization of an EGFP-LSm4 fusion protein in living cells following DNA damage induction via laser-microirradiation. EGFP-LSm4 exhibits rapid and transient accumulation at microirradiated tracks, appearing within 15 seconds and persisting for over 15 minutes (Figure 1E). Intriguingly, LSm1-3 and LSm5-8 proteins, which do not form BCs, were not recruited to laser-microirradiated sites (Supplementary Figure 1D), suggesting that BC forming capacity correlates with the recruitment to DNA damage site. To identify the region of LSm4 responsible for its accumulation at damage sites, we performed deletion mapping analysis (Figure 1F). An LSm4 mutant lacking the C-terminal region spanning amino acids 57- 139 (LSm4^ΔC^) failed to accumulate at laser-microirradiated lesions. Further deletion analysis pinpointed residues 69-92 as critical for LSm4 recruitment, as the deletion mutant LSm4^Δ69–92^ showed no accumulation at DNA damage sites (Figure 1G, Supplementary Figure 1E). Interestingly, we noticed that LSm4^Δ69–92^ also lost its ability to form condensates at the nucleolus (Figure 1G). These results prompted us to test whether this mutant retains the capacity to undergo LLPS and forms BCs. Using OptoDroplet assay, we found that while LSm4-Cry2WT forms robust light-induced OptoDroplet foci, LSm4^Δ69–92^ mutant completely lost its ability to form OptoDroplets (Figure 1H, Supplementary Movie 5). Collectively, these data demonstrate that residues 69-92 of LSm4 are required both for its LLPS capacity and for its accumulation at DNA damage sites, supporting a model in which LSm4 is recruited to DNA damage via LLPS-driven condensation. Moreover, our findings identify the LSm4 region spanning residues 69-92 as the critical determinant that functionally distinguishes LSm4 from other LSm proteins. Consistent with this model, sequence conservation analysis revealed that 69–92 region of LSm4 is highly conserved in LSm4 protein across species, yet absent from other LSm family members (Figures 1I, 1J), underscoring its evolutionary importance and LSm4-specific function.

### LSm4 forms biomolecular condensates at DSBs in transcriptionally active regions

The identification of LSm4 within the interactome of key DSB repair proteins ^55^ prompted us to investigate whether it is recruited to DSB sites. Herein, we present multiple lines of evidence demonstrating that LSm4 is specifically recruited to DSBs occurring in transcriptionally active chromatin. Biochemical fractionation showed that LSm4 accumulates on chromatin following treatment with the DSB-inducing agent Zeocin (Figure 2A). Given that LSm4 is an RNA-binding protein, we asked whether RNA molecules contribute to LSm4 accumulation at DSBs. Remarkably, RNase A treatment abolished LSm4 enrichment at damaged chromatin, suggesting that the presence of RNA molecules facilitates LSm4 recruitment to chromatin following DNA damage (Figure 2B). To directly test LSm4 recruitment to known DSBs, we utilized DIvA (DSB Inducible via *Asi*SI) U2OS cells system to induce DSBs at annotated *Asi*SI sites, followed by chromatin immunoprecipitation and sequencing (ChIP-seq). LSm4 ChIP-seq analysis revealed that it accumulates predominantly at DSBs localized within transcriptionally active regions, while its recruitment to DSBs in silent chromatin was markedly reduced (Figure 2C). Next, we performed ChIP followed by quantitative real-time PCR (ChIP-qPCR) and confirmed LSm4 accumulation at three individual *Asi*SI sites adjacent to the actively transcribed genes *MIS12*, *RBMXL1*, and *KLHL22* (Figures 2D, 2E).

**Figure 2:**
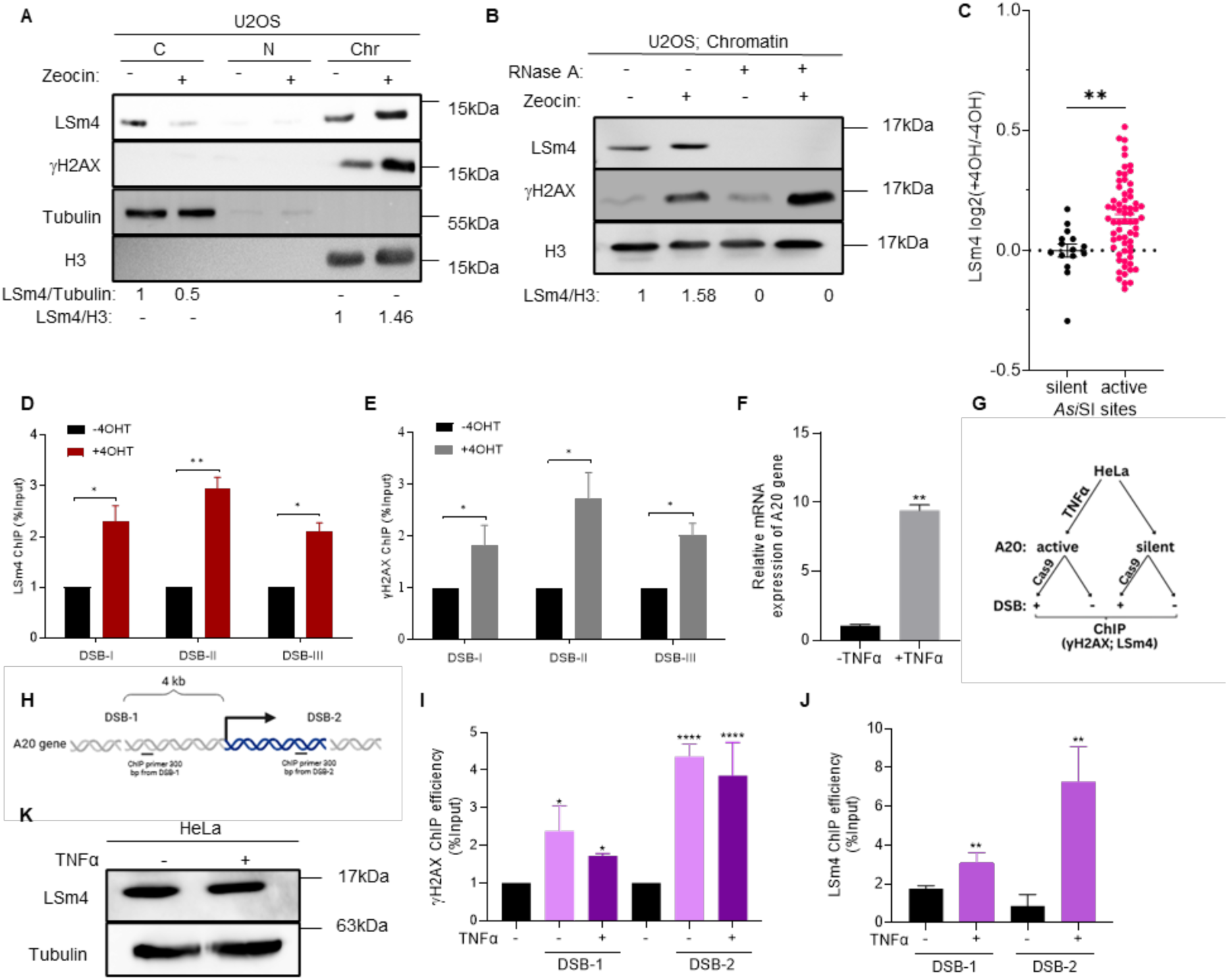
LSm4 forms biomolecular condensates at DSBs in transcriptionally active regions. (A) Immunoblot analysis of LSm4 subcellular localization in control and Zeocin-treated U2OS cells. C: cytoplasmic fraction; N: nuclear soluble fraction; Chr: chromatin-bound fraction. (B) Immunoblot analysis of LSm4 chromatin enrichment in control and Zeocin-treated U2OS cells upon RNase A treatment. (C) LSm4 ChIP-seq enrichment in 4-OHT treated and untreated U2OS-DIvA cells at transcriptionally active or silent/intergenic DSBs. Each dot corresponds to the average log2 -/+4OHT value at a single *Asi*SI site. Statistical significance was determined by nonparametric unpaired Mann-Whitney-Wilcoxon test (P = 0.0034). (D and E) ChIP-qPCR for LSm4 (D) and γH2AX (E) in U2OS-DIvA cells before and after 4-OHT treatment. Primers used to capture *Asi*SI sites located nearby the transcriptionally active genes *RBMXL1* (DSB-I), *MIS12* (DSB-II), and *KLHL22* (DSB-III). (F) Relative mRNA expression of *A20* gene in HeLa cells upon treatment with TNFα. GAPDH is used as control. (G) Schematic representation of ChIP-qPCR experiment to measure LSm4 recruitment to CRISPR- Cas9-induced DSBs at active and inactive *A20* gene. (H) Schematic diagram visualizing the position of two DSBs induced by Cas9 at the *A20* gene, and the position of RT-PCR primers used to test LSm4 and γH2AX enrichment in HeLa cells. (I and J) ChIP-qPCR for γH2AX (I) and LSm4 (J) at DSBs nearby *A20* gene in HeLa cells. Error bars represent as mean ± SD; n=3. P value was determined by two-sided Students t-test relative to control cells. ns is not significant, *p<0.05, ∗∗p<0.01, ***p<0.001, and ****p<0.001. (K) Immunoblot analysis of LSm4 levels in control and TNFα-treated HeLa cells, representative of n=3.

To directly assess how transcriptional state governs LSm4 recruitment at a defined genomic locus, we leveraged the elegant A20 reporter system that we had previously established ^24^. *A20* gene is transcriptionally silent in HeLa cells but can be rapidly activated by treatment with Tumor Necrosis Factor-alpha (TNF-α), as evident by ∼ten-fold increase in its mRNA levels (Figures 2F). We introduced Cas9-mediated DSBs at two different regions within the *A20* locus both in its silent and transcriptionally active states, followed by ChIP-qPCR for γH2AX and LSm4 (Figure 2G, 2H). Results showed that while γH2AX is comparably induced upon DSB induction at both active and silent *A20* loci, LSm4 exhibited significantly greater enrichment at the DSB sites when *A20* was transcriptionally active (Figures 2I, 2J). Importantly, the accumulation of LSm4 at DSBs within *A20* gene following TNF-α treatment is not due to an increase in total LSm4 protein levels (Figure 2K). Collectively, these data demonstrate that LSm4 condenses preferentially at DSBs within transcriptionally active chromatin, suggesting that transcriptional status dictates LSm4 condensation at damage sites.

### LSm4 drives RNA decay at DSBs

Previous work demonstrated that cytoplasmic LSm1-7 complex mediates RNA turnover at P-bodies condensates in yeast ^65^. Therefore, we wondered whether the DNA damage-induced BCs of LSm4 promote RNA decay at DSB sites. To address this, we employed two orthogonal and complementary assays to monitor the RNA decay kinetics of transcriptionally active genes located in close genomic proximity to site-specific *Asi*SI-induced DSBs in DIvA cells. First, we employed a transcription shutoff assay using DRB to quantitatively measure transcript stability upon DSB induction ^66^. LSm4-proficient and deficient cells were treated with 4- hydroxytamoxifen (4-OHT) to induce DSBs at *Asi*SI sites, then, transcription was globally inhibited with DRB and RNA was harvested at 1, 4, and 6 hours post-inhibition (Figure 3A). Reverse transcription-qPCR (RT-qPCR) analysis revealed that depletion of LSm4 significantly attenuated the decay of MIS12 and AGA nascent transcripts following DSB induction (Figures 3B, 3C), indicating that LSm4 is required for efficient RNA clearance at DSB sites. Second, we measured RNA decay using Roadblock-qPCR assay. Unlike the aforementioned transcription shutoff assay, the Roadblock method circumvents the global perturbations of transcription inhibition by using a brief pulse of 4-thiouridine (4sU) to label newly transcribed RNAs, followed by N-ethylmaleimide (NEM) treatment to chemically block reverse transcription of these labeled transcripts. This allows for the specific quantification of the pre-existing, non-labeled endogenous RNA pool over time ^67,68^. We applied this assay in DIvA cells following induction of *Asi*SI-mediated DSBs at *RBMXL1* and *MIS12* genes (Figure 3D). Results showed that LSm4-proficient cells exhibited a rapid decline in the pre-existing mRNA levels of *RBMXL1* and *MIS12* genes, which reflects the active RNA decay of those transcripts. Intriguingly, RNA decay was markedly attenuated in LSm4-deficient cells, reflecting a significant stabilization of the pre-existing mRNAs (Figures 3E and 3F). Altogether, these data establish that LSm4 is required for efficient decay of RNA transcribed from genes proximal to DSBs.

**Figure 3:**
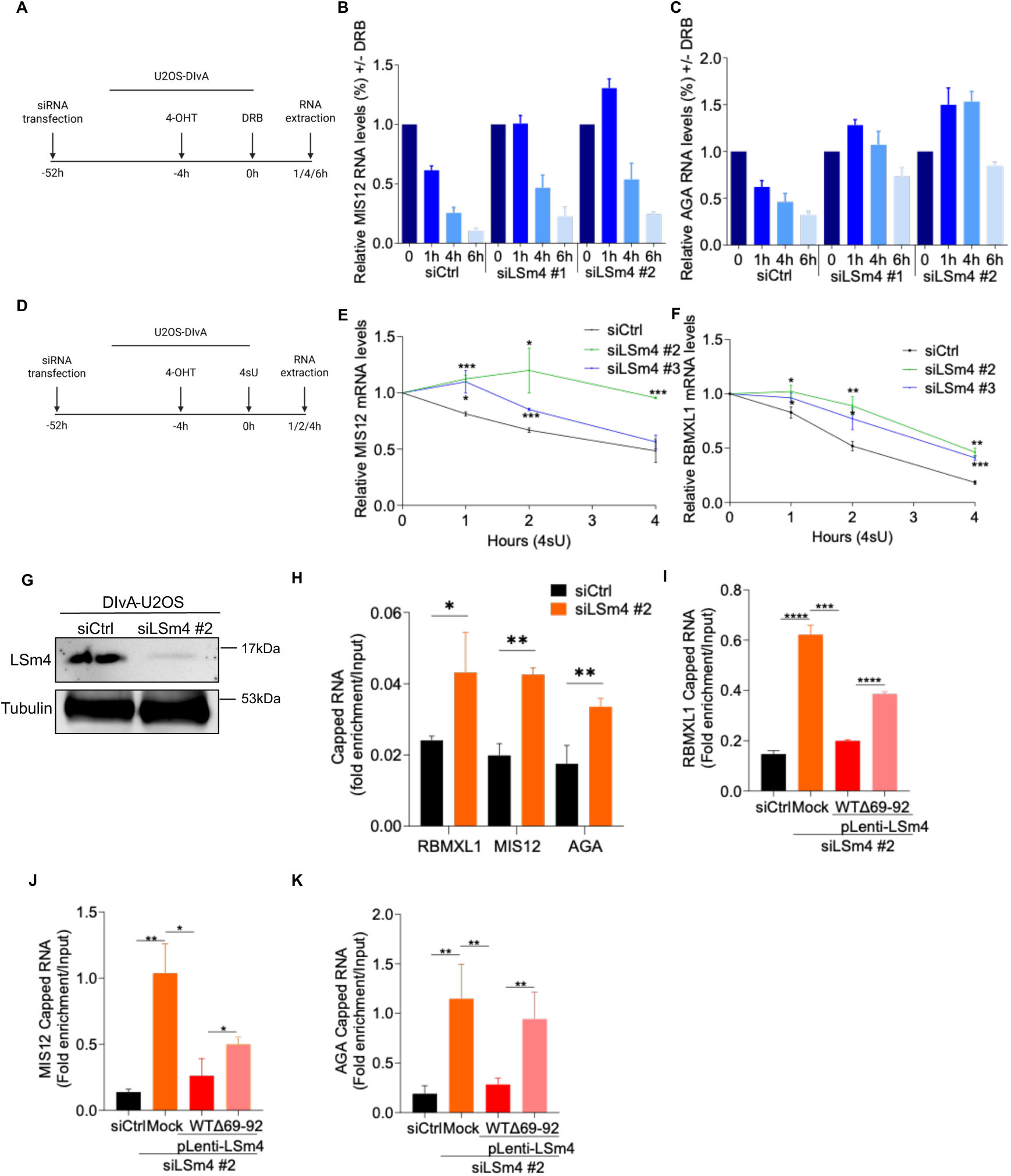
LSm4 biomolecular condensates promote RNA decapping to drive RNA decay at DSBs. (A) Schematic diagram of transcription shutoff assay using DRB. (B and C) mRNA decay analysis of MIS12 (B) and AGA (C) transcripts using transcriptional shutoff assay in U2OS-DIvA cells following DSB induction. (D) Schematic diagram of Roadblock-qPCR assay. (E and F) mRNA decay analysis of MIS12 (E) and RBMXL1 (F) transcripts using Roadblock-qPCR assay in U2OS-DIvA cells following DSB induction. (G) Immunoblot analysis of LSm4 levels in U2OS-DIvA cells treated with siCtrl or siLSm4 #2. (H) Cap-IP for RBMXL1, MIS12, and AGA transcripts following DSB induction in U2OS-DIvA cells. (I-K) Cap-IP for RBMXL1 (I), MIS12 (J) and AGA (K) transcripts in control and siLSm4-depleted cells expressing LSm4^WT^ or LSm4^Δ69-92^ mutant following DSB induction in U2OS-DIvA cells. All data are presented as mean ±SD; n=3. P value was determined by two-sided Students t-test relative to control cells. ns is not significant, *p<0.05, ∗∗p<0.01, ***p<0.001, and ****p<0.001.

### LSm4 biomolecular condensates promotes RNA decapping to drive RNA decay at DSBs

Given that the cytoplasmic LSm1-7 complex promotes RNA decapping to initiate 5’→3’ decay, we hypothesized that LSm4 facilitate a similar decapping for chromatin-associated RNA at DNA damage sites. To test this, we performed cap immunoprecipitation (Cap-IP) on chromatin-associated RNA from DIvA cells using an anti-m7G cap antibody. Results demonstrated that upon DSB induction, the levels of capped MIS12, RBMXL1, and AGA1 transcripts were significantly elevated in LSm4-deficient DIvA cells. (Figures 3G, 3H). We concluded therefore that LSm4 underpins decapping of nuclear RNA adjacent to DSB sites, plausibly to promote the efficient clearance of nascent RNA from damage sites.

To determine whether the role of LSm4 in promoting decapping is dependent on its ability to form damage-induced BCs, we performed Cap-IP in LSm4-deficient cells stably expressing either LSm4^WT^ or LSm4^Δ69–92^ mutant, which lost its ability to form BCs at DSBs. Results showed that expression of LSm4^WT^ fully restored efficient decapping, whereas the LSm4^Δ69–92^ mutant did not (Figures 3I-K). This indicates that LSm4 BCs are critical for promoting RNA decapping at DSBs. Collectively, our results establish that LSm4 forms BCs at DSBs to facilitate RNA decapping, thereby promoting efficient clearance of nascent RNA from damage sites.

### LSm4 regulates RNA decapping and decay independently of its roles in transcription and splicing

To investigate whether LSm4-mediated RNA decay and decapping at DSBs is influenced by its canonical roles in gene expression or pre-mRNA splicing, we performed global transcriptomic profiling. We conducted RNA sequencing (RNA-seq) in U2OS cells following depletion of LSm4 using two independent siRNAs (Figure 4A). Bioinformatic analysis identified 1,052 genes with significantly altered expression levels upon LSm4 knockdown (Figure 4B, Supplementary Figures 2A, 2B, Supplementary Table S1). Gene Ontology (GO) analysis revealed that these differentially expressed (DE) genes were enriched in several pathways, including those associated with oncogenic signaling (Figure 4C). Crucially, the expression levels of core RNA decapping and decay factors remained unaffected by LSm4 depletion. Given the established role of the nuclear LSm2-8 complex in U6 snRNA stability, we next performed a genome-wide splicing analysis. We identified thousands of differential alternative splicing (AS) events affecting 461 genes, with a predominant shift toward exon inclusion (Figure 4D, Supplementary Figure 2C, Supplementary Table S2). Among the 1,052 DE genes, 223 displayed significant alternative splicing events, predominantly exon skipping (Figure 4E, Supplementary Figure 2D). Notably, the splicing patterns of established RNA decapping or decay genes were not altered in LSm4-deficient cells.

**Figure 4:**
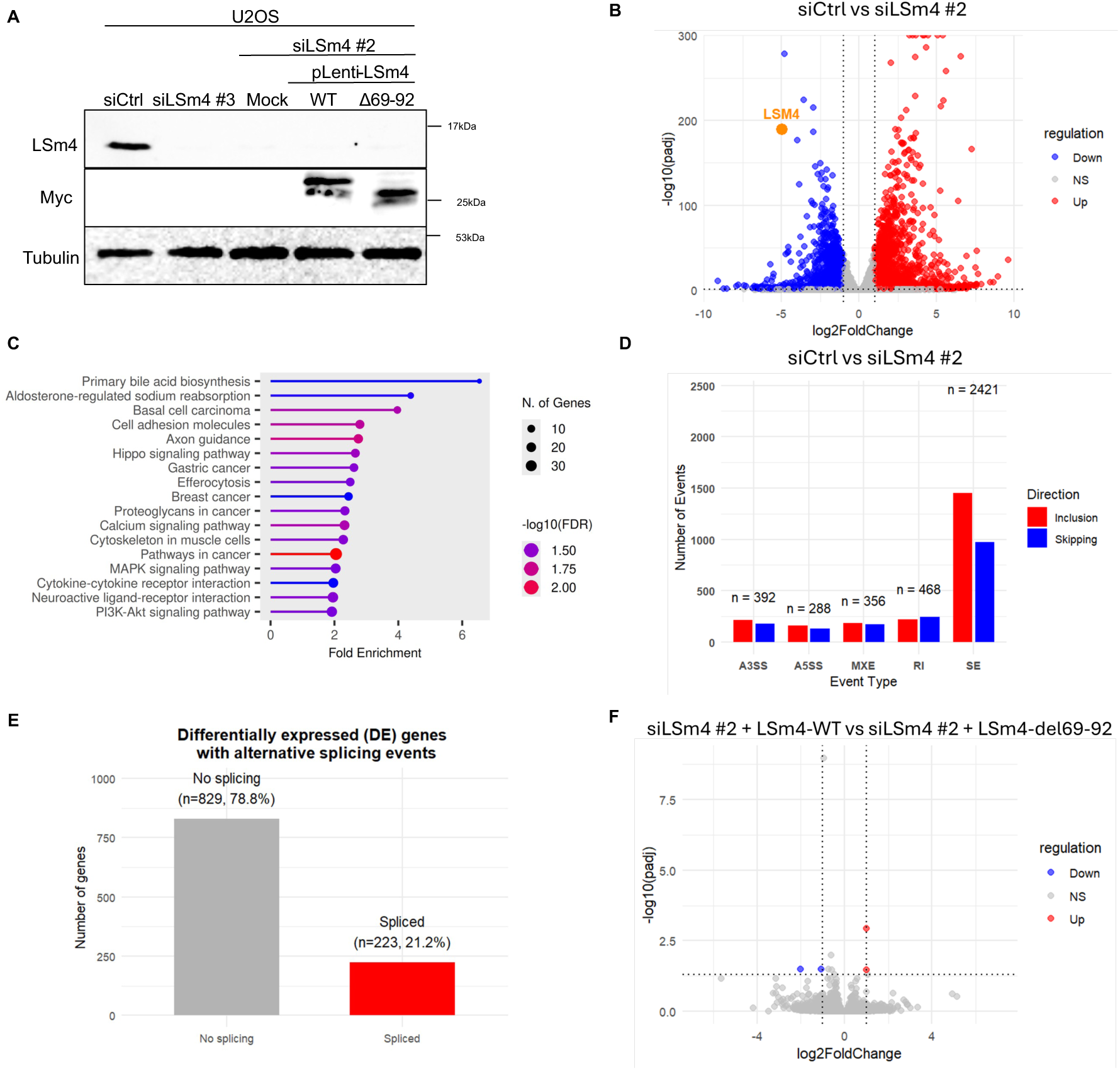
LSm4 regulates RNA decapping and decay independently of its roles in transcription and splicing. (A) Immunoblot analysis of LSm4 and Myc in samples sent for RNA-seq. (B) Volcano plot summarizing differential gene expression data obtained from RNA-seq analysis between siCtrl- and siLSm4 #2-transfected U2OS cells. Upregulated genes with log2FoldChange(siLSm4 #2/control) > 1 and *P*adj-value < 0.05 are marked in red (n=2333), while downregulated genes with log2FoldChange(siLSm4 #2/control) < -1 and *P*adj-value < 0.05 are marked in blue (n=2204). (C) KEGG pathway enrichment analysis of differentially expressed (DE) genes identified by RNA-seq and shared between both siLSm4-treated samples relative to control U2OS cells. (D) Summary of significant alternative splicing events leading to skipping (blue) or inclusion (red) observed upon siLSm4 #2 treatment as detected by rMATS. Significantly altered splicing events were classified as having a minimum inclusion level difference of 0.1, *p* value < 0.01, and FDR < 0.05. SE: skipped exon, MXE: mutually exclusive exons, A5SS: alternative 5’ splice site, A3SS: alternative 3’ splice site, IR: intron retention. (E) Bar plot indicating the number of DE genes with significantly altered splicing events identified by RNA-seq and shared between both siLSm4- treated samples relative to control U2OS cells. (F) Volcano plot summarizing differential gene expression data obtained from RNA-seq analysis between siLSm4 #2 cells expressing LSm4^WT^ and LSm4^Δ69-92^ U2OS cells. Upregulated genes with log2FoldChange(siLSm4 #2 + LSm4^Δ69-92^/ siLSm4 #2 + LSm4^WT^) > 1 and *P*adj-value < 0.05 are marked in red, while downregulated genes with log2FoldChange(siLSm4 #2 + LSm4^Δ69-92^/ siLSm4 #2 + LSm4^WT^) < -1 and *P*adj- value < 0.05 are marked in blue.

To further substantiate that the observed defects in RNA clearance at DSBs are specifically linked to LSm4 biomolecular condensation, we performed RNA-seq in LSm4-deficient cells complemented with either LSm4^WT^ or the LSm4 ^Δ69–92^ mutant (Figure 4A). Strikingly, a comparison of these transcriptomes revealed only four genes whose expression was significantly altered between the two conditions (Figure 4F, Supplementary Figure 2E). None of these genes are involved in RNA decapping or decay pathways. Collectively, these results demonstrate that LSm4 regulates localized RNA clearance through its intrinsic capacity for biomolecular condensation at DSBs, rather than by modulating the global transcriptome or splicing machinery.

### LSm4 recruits XRN2 to DSBs to drive RNA decay

To shed molecular insight into how LSm4 drives RNA decay at DSBs, we sought to identify the ribonuclease responsible for executing RNA degradation. We hypothesized that XRN2 acts downstream of LSm4 to clear transcripts at damage sites, since it has an established role as a nuclear 5’→3’ exoribonuclease. To address this hypothesis, we first asked whether XRN2 promotes RNA decay at DSB sites using the Roadblock-qPCR assay. Indeed, knockdown of XRN2 led to significant stabilization of pre-existing RBMXL1 transcripts following DSB induction (Figures 5A-B), phenocopying the defect observed upon LSm4 depletion (Figures 3E, 3F). We next examined whether XRN2 is recruited to DSB sites. Laser-microirradiation assay revealed rapid recruitment of endogenous XRN2 to laser-induced DNA damage sites, which were co-localized with γH2AX stripes (Figure 5C). In addition, immunofluorescence analysis showed a pronounced increase in the chromatin-associated signal of XRN2 following ionizing radiation (IR) (Figure 5D). Moreover, ChIP-qPCR in DIvA cells showed enrichment of XRN2 at *Asi*SI-induced DSBs at actively transcribed chromatin (Figure 5E). Together, these three complementary approaches establish XRN2 as a bona fide component of the DSB response. Next, we tested whether LSm4 regulates XRN2 recruitment. Immunofluorescence analysis revealed that LSm4 depletion severely impairs the IR-induced accumulation of endogenous XRN2 on chromatin, while not affecting XRN2 protein levels (Figures 5F, Supplementary Figure 3A). Consistently, ChIP-qPCR analysis in DIvA cells revealed that LSm4 depletion significantly impaired XRN2 accumulation at *Asi*SI-induced DSBs (Figure 5G). To determine if XRN2 accumulation at DSB sites is dependent on the phase-separation capacity of LSm4, we complemented LSm4-deficient cells with either LSm4^WT^ or the LLPS-deficient mutant LSm4^Δ69–92^. While LSm4^WT^ fully restored XRN2 recruitment to DSBs, LSm4^Δ69–92^ mutant failed to do so (Figures 5H, 5I), indicating that LSm4 condensates are critical for XRN2 accumulation at damage sites. Collectively, these findings delineate a pathway in which LSm4 forms LLPS condensates at DSBs to recruit XRN2 exonuclease, thereby enabling the efficient RNA decay at DSB microenvironment.

**Figure 5:**
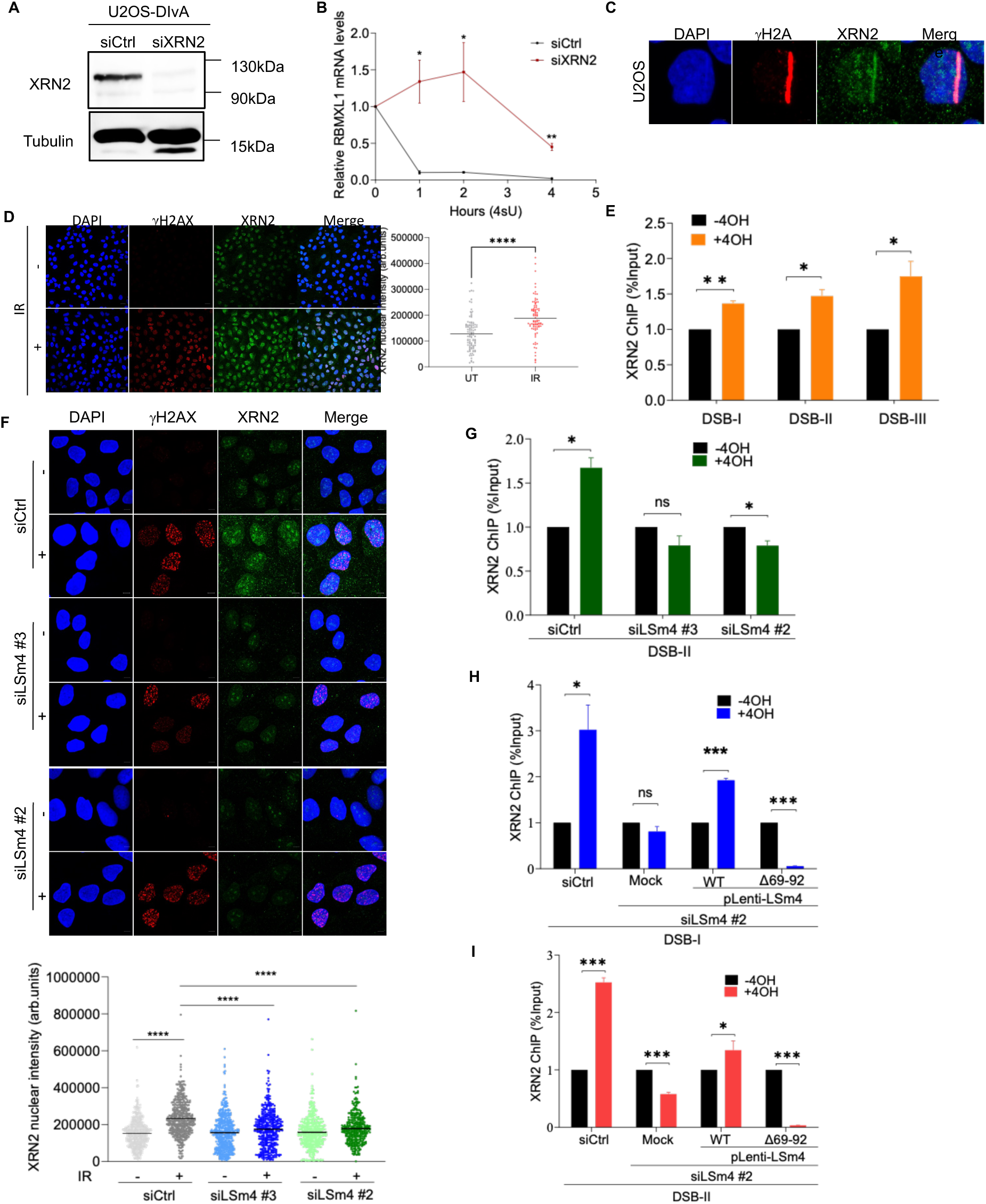
XRN2 is recruited to DSBs by LSm4 to facilitate RNA decay. (A) Immunoblot analysis for XRN2 levels in control and siXRN2-depleted U2OS cells. (B) mRNA decay analysis of RBMXL1 transcript using Roadblock-qPCR assay in DIvA cells following DSB induction. Data are presented as mean ± SD; n=3. P value was determined by two-sided Students t-test relative to control cells. *p<0.05 and ∗∗p<0.01. (C) Representative U2OS cell showing endogenous XRN2 (green) accumulation at laser-microirradiated regions 10 minutes after damage induction. γH2AX (red) marks DNA damage sites, and DNA is stained with DAPI (blue). (D) Left: representative image of immunofluorescence assay measuring XRN2 chromatin association before and after 4Gys IR. Right: quantification of XRN2 signal on chromatin in U2OS cells before and after IR. Horizontal bars represent mean intensity per nucleus ±SEM. UT: n=104; IR: n=90. P value was determined by two-tailed Mann–Whitney test. ∗∗∗∗p<0.0001. (E) ChIP-qPCR for XRN2 in U2OS-DIvA cells before and after 4-OHT treatment. Data are presented as mean ±SD; n=3. (F) Top: representative image of immunofluorescence assay measuring XRN2 chromatin association in LSm4-proficient and deficient cells, before and after 4Gys IR. Bottom: quantification of XRN2 signal. Horizontal bars represent mean intensity per nucleus ±SEM; n>350. P value was determined by two-tailed Mann–Whitney test. ∗∗∗∗p<0.0001. (G) As in (E) except in LSm4-proficient and deficient U2OS-DIvA cells. (H and I) As in (E) except in LSm4-proficient and deficient U2OS-DIvA cells expressing either LSm4^WT^ or LSm4^Δ69–92^ in *MIS12* (H) and *RBMXL1* (H) genes. Data are presented as mean ±SD; n=3. P value was determined two-sided Students t-test relative to control cells. ns is not significant, *p<0.05, ∗∗p<0.01 and ***p<0.001.

### LSm4 regulates R-loop homeostasis at DSB sites

Having established that LSm4 drives the decay of nascent RNA at DSBs, we reasoned that its loss might lead to the persistence of R-loops, which are known to impede repair and promote genomic instability ^34,35^. We first assessed the global R-loop levels using dot-blot assay. Results showed that LSm4 depletion resulted in a significant increase in R-loop levels (Figure 6A). This finding was corroborated by immunofluorescence analysis using a purified GFP-tagged catalytically inactive mutant of RNase H1 (GFP-dRNH1), which binds R-loops and serves as a fluorescent reporter for their abundance ^69^ (Figure 6B). To determine whether this effect occurs specifically at DSBs, we performed DNA:RNA immunoprecipitation followed by qPCR (DRIP-qPCR) in DIvA cells. The specificity of the DRIP-qPCR assay was validated by measuring R-loop levels at two control loci: *SNRNP* (R-loop-poor) and *RPL13A* (R-loop-enriched) (Supplementary Figure 4A). Following 4-OHT treatment, we observed a significant increase in R-loops at transcriptionally active genes proximal to *Asi*SI-induced DSBs upon LSm4 depletion (Figure 6C). Notably, persistent R-loops are known to impair the formation of RAD51 foci ^30,33,35–41^. We therefore asked whether elevated R-loops in LSm4-deficient cells compromise RAD51 recruitment. Immunofluorescence analysis revealed that LSm4 depletion significantly impaired the formation of RAD51 foci following IR (Figure 6D-E). This defect is not due to reduced RAD51 RNA and protein levels in LSm4-deficient cells (Supplementary Figure 4B, Supplementary Table 1). Consistently, RAD51 ChIP-qPCR in DIvA cells showed that LSm4 loss reduces RAD51 recruitment to *Asi*SI-induced DSBs (Figure 6F). To directly test whether the RAD51 recruitment defect is a consequence of local R-loop accumulation rather than an indirect effect of LSm4 loss, we sought to resolve R-loops specifically at *Asi*SI- induced DSB site nearby *MIS12* gene. We targeted a catalytically dead Cas9 (dCas9)-fused to RNase H1 domain to the *MIS12* locus (Figure 6G, Supplementary Figure 4C). Strikingly, this site-specific R-loop depletion in LSm4-deficient cells was sufficient to restore RAD51 recruitment to the DSB (Figure 6H). Collectively, these data demonstrate that LSm4-mediated RNA clearance maintains R-loop homeostasis at DSBs, thereby facilitating the critical step of RAD51 recruitment plausibly to underpin homologous recombination.

**Figure 6:**
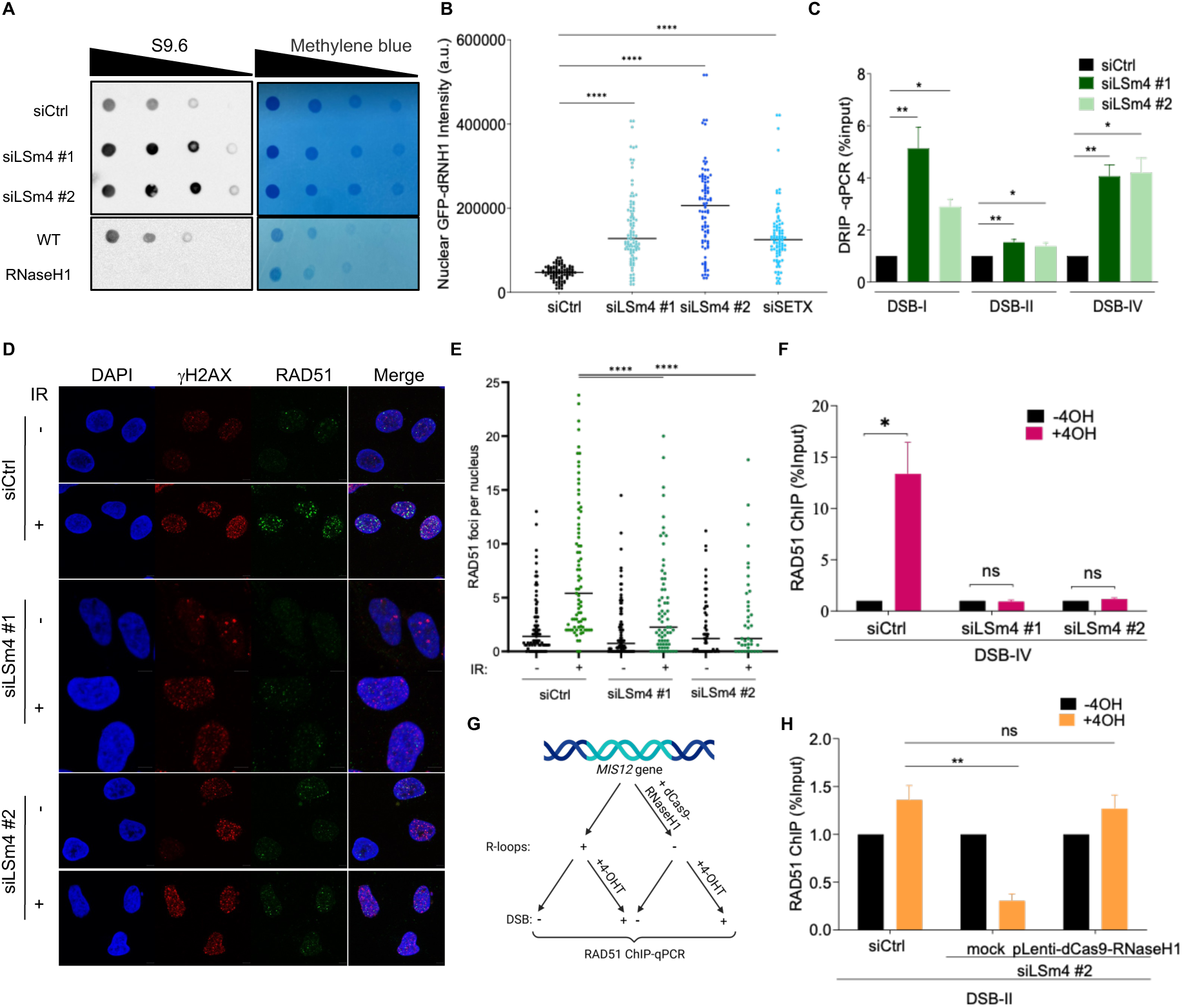
LSm4 regulates R-loop homeostasis and facilitates RAD51 recruitment at DSB sites. (A) Dot-blot analysis in LSm4-proficient and deficient U2OS cells. Top: Methylene blue staining as loading control. Bottom: S9.6 antibody staining. RNase H1 serves as negative control. (B) Quantification of nuclear mean intensity of purified catalytically inactive GFP- dRNH1 in U2OS cells transfected with indicated siRNAs. Depletion of SETX serves as positive control. Horizontal bars represent mean intensity per nucleus ±SEM; n>85. P value was determined by two-tailed Mann–Whitney test. (C) DRIP-qPCR in LSm4-proficient and deficient U2OS-DIvA cells following 4-OHT treatment at DSBs nearby transcriptionally active genes. Data are presented as mean ±SD; n=3. P value was determined by two-sided Students t-test relative to control cells. (D) Representative immunofluorescence images of RAD51 and γH2AX foci in LSm4-proficient and deficient U2OS cells before and after 4Gys IR. DNA is stained with DAPI. (E) Quantification of (D). Horizontal bars represent mean foci number per nucleus ±SEM; n>200. P value was determined by two-tailed Mann–Whitney test. (F) RAD51 ChIP-qPCR in LSm4-proficient and deficient U2OS-DIvA cells. (G) Schematic diagram of dCas9-RNaseH1 tethering experiment. (H) Same in (F) except in LSm4-proficient and deficient U2OS-DIvA cells infected with either mock or dCas9-RNaseH1-HA. Data are presented as mean ±SD; n=3. P value was determined by two-sided Students t-test relative to control cells. ns is not significant, *p<0.05, ∗∗p<0.01, ***p<0.001 and ****p<0.0001.

### LSm4 condensates are essential for homologous recombination repair

Our data establish that LSm4 condensates facilitate RNA clearance and prevent R-loop accumulation to enable RAD51 recruitment to DSBs. We therefore asked whether loss of LSm4 leads to HR deficiency. Initially, we showed that LSm4 depletion resulted in elevated baseline levels of γH2AX, as shown by both western blot and immunofluorescence analysis (Figures 7-B). Interestingly, γH2AX signal in LSm4 deficient cells was further exacerbated following DNA damage, pointing to defective DSB repair. To directly measure HR efficiency, we employed mClover-based reporter system, which quantifies HR repair of a Cas9-induced DSB upstream of the LMNA gene via a gene conversion event that restores a functional fluorescent mClover-LMNA protein ^70^. Results showed that LSm4 depletion significantly impaired HR, as evidenced by a marked reduction in mClover-positive cells (Figure 7C, Supplementary Figures 5A-B). Importantly, our transcriptomic data confirmed that LSm4 depletion does not alter the expression of core DSB repair factors (Supplementary Table 1), suggesting that LSm4 directly promotes HR repair independent of gene expression changes. To assess the impact of LSm4 deficiency on genomic stability, we monitored the formation of DNA translocations, a hallmark of defective HR, following DSB induction in DIvA cells. LSm4 depletion significantly increased translocation frequency, similar to the phenotype caused by loss of the R-loops helicase SETX (Figure 7D). Therefore, we concluded that LSm4 is essential for suppressing illegitimate rejoining and preserving genomic integrity. Finally, to determine if the HR repair depends on LSm4 phase-separation capacity, we complemented LSm4-deficient cells with either LSm4^WT^ or the LLPS-deficient mutant LSm4^Δ69–92^. While re-expression of LSm4^WT^ restored HR efficiency in the mClover reporter assay, the LSm4^Δ69–92^ failed to rescue the defective HR repair (Figure 7E, Supplementary Figure 5C). Altogether, we concluded that LSm4 LLPS condensates drive RNA clearance to suppress R-loops, enabling RAD51 loading and thereby ensuring intact HR.

**Figure 7:**
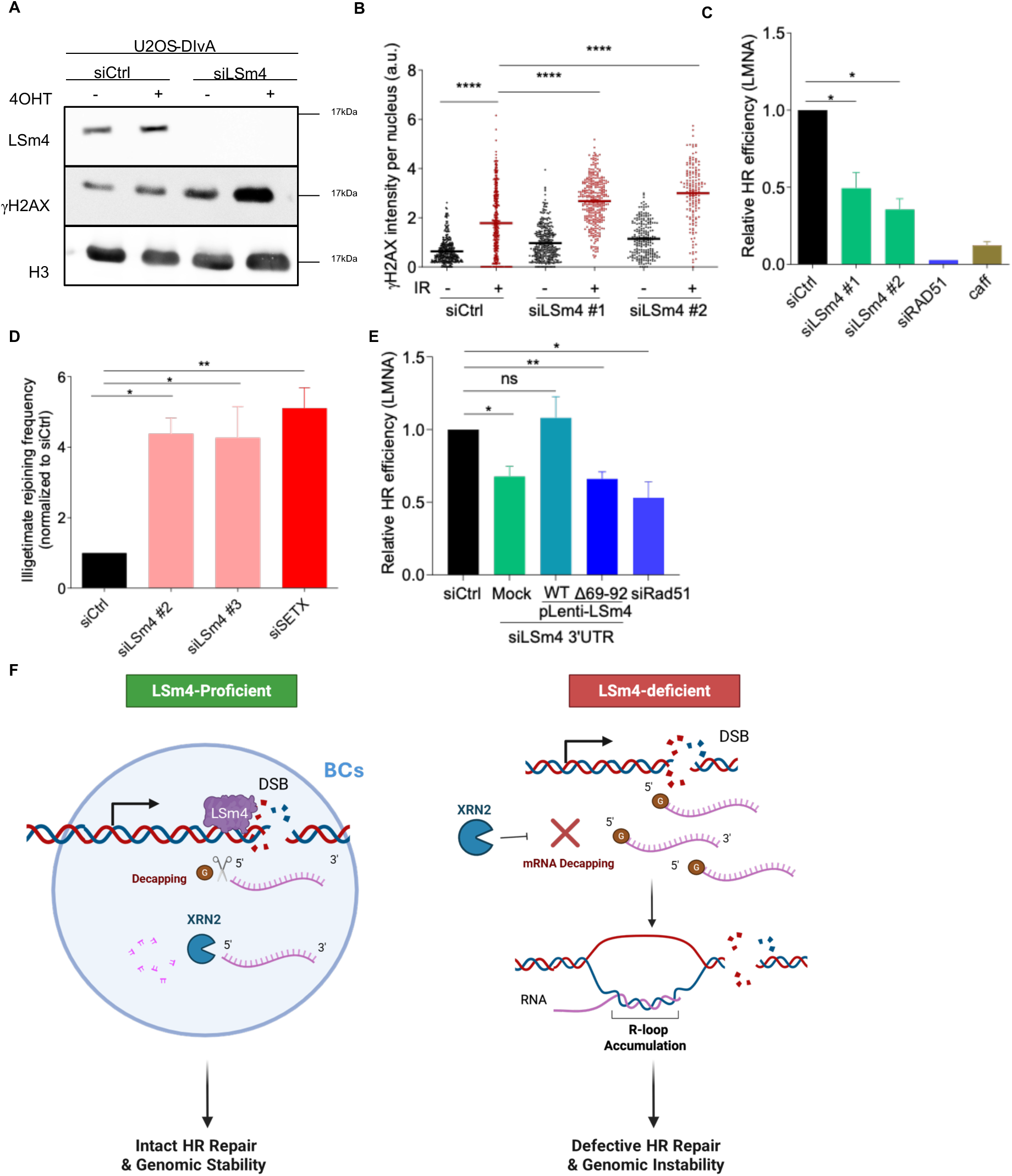
LSm4 condensates are essential for homologous recombination repair and genomic stability. (A) Immunoblot analysis for LSm4 and γH2AX levels before and after DSB induction in U2OS-DIvA cells. H3 serves as loading control. (B) γH2AX mean intensity per nucleus in LSm4-proficient and deficient U2OS cells before and after 4Gy IR. Horizontal bars represent mean foci number per nucleus ±SEM; n>150. P value was determined by two-tailed Mann–Whitney test. (C) LSm4 depletion impairs HR of endogenous DSBs induced by Cas9 endonuclease upstream the LMNA gene. LSm4-proficient and deficient U2OS cells were transfected with fluorescence (clover)-based reporter, and HR efficiency was determined as described in the Methods section. Error bars represent standard deviation from the mean. RAD51 depletion and Caffeine (caff) were used as controls. (D) Illegitimate rejoining frequency between two transcription-coupled DSBs (MIS12 and LYRM2; chr17:5,390,209 and chr6:9,034,817) in U2OS-DIvA cells following DSB induction and repair (+4-OHT, +IAA). All samples are normalized to siCtrl. Data are presented as mean ±SD; n=3. (E) LSm4 BCs are required for efficient HR repair of DSBs. HR efficiency in U2OS was determined as described in (C). (F) Model summarizing the role of LSm4 in RNA clearance and repair. Briefly, damage-induced LSm4 BCs promote RNA decapping and XRN2-mediated RNA decay at DNA DSBs, thereby facilitating efficient HR repair and preserving genomic stability. ns is not significant, *p<0.05, ∗∗p<0.01, ***p<0.001, ****p<0.0001.

## Discussion

Our study unveils a previously unrecognized role for LSm4 in orchestrating localized RNA decay at DNA DSBs through biomolecular condensate (BC) formation. We demonstrate that LSm4, but not other LSm1-8 proteins, undergoes liquid-liquid phase separation (LLPS) to form dynamic condensates specifically at DSBs within transcriptionally active chromatin. These damage-induced BCs serve as specialized microenvironments that concentrate the RNA decay machinery, XRN2 exonuclease and presumably decapping enzymes, thereby facilitating rapid degradation of nascent transcripts proximal to break sites. This LSm4-driven RNA clearance mechanism is essential for suppressing pathological R-loop accumulation, enabling efficient RAD51 filament assembly, and ensuring high-fidelity HR repair (Figure 7F).

The LSm protein family has been extensively studied for its roles in RNA metabolism, where the cytoplasmic LSm1-7 complex mediates mRNA decay and the nuclear LSm2-8 complex functioning in pre-mRNA splicing ^44,45,50,71–73^. Our results showing that only LSm4, among all LSm1-8 subunits, forms BCs at DSBs reveals an unexpected distinct function of LSm4. This observation is consistent with the recent identification of LSm4 as a phase-separating protein in a high-throughput endogenous proteomic screen ^74^ and with the ability of recombinant LSm4 protein to undergo LLPS *in vitro* ^57^. To identify the sequence that distinguishes LSm4 from its paralogs, we mapped residues 69-92 as the critical determinant of the phase separation capacity of LSm4. By virtue of this unique domain, LSm4 acquires a damage-inducible capacity to orchestrate nuclear RNA clearance at DSB sites, representing a regulatory mechanism that is functionally partitioned from its canonical roles in splicing and cytoplasmic decay.

The observation that LSm4 condensates preferentially at DSBs in transcriptionally active regions, but not in silent chromatin, suggests that LSm4 recruitment is coupled to the presence of nascent RNA transcripts. Indeed, our finding that RNase A treatment abolishes LSm4 chromatin enrichment following DNA damage supports the notion that LSm4 is recruited to DSBs in an RNA-dependent manner. This transcription-coupled recruitment may represent an elegant mechanism for resolving the transcription-repair conflict, whereby LSm4 BCs concentrate the decay machinery precisely at sites where transcription and DNA damage intersect.

Our findings contribute to the emerging paradigm that BCs function as organizational hubs that enhance the efficiency and specificity of the DNA damage response. Several RNA-binding proteins, including FUS, EWSR1, and TAF15, have been shown to undergo PARP1-dependent phase separation at DNA damage sites, where they facilitate DNA end resection and recruit downstream repair factors ^64,75^. Similarly, 53BP1 forms liquid-like condensates that promote non-homologous end joining, while proteins such as RAP80 concentrate at damage sites through ubiquitin-dependent phase separation mechanisms ^76,77^. LSm4 BCs at DSBs appear to serve a fundamentally different yet complementary function by orchestrating the removal of RNA obstacles that would otherwise impede repair, rather than directly participating in DNA end processing or repair pathway choice. This functional distinction highlights how phase separation can be exploited for diverse purposes within the damage response. Some condensates sequester repair proteins to enhance local concentration and activity, while LSm4 condensates create specialized zones for RNA clearance.

The recruitment of XRN2 to DSBs by LSm4 condensates establishes a direct mechanistic link between RNA decay and HR. XRN2 is a well-characterized nuclear 5′→3′ exoribonuclease with established roles in transcription termination, processing of small nuclear RNAs, and degradation of aberrant transcripts. Our work reveals an additional layer of XRN2 function, through the targeted elimination of nascent transcripts at DSBs to prevent R-loop accumulation. This function is critically dependent on LSm4, as XRN2 recruitment to damage sites is severely impaired in LSm4-deficient cells and cannot be rescued by the LLPS-deficient LSm4^Δ69–92^ mutant. Our findings are consistent with previous studies showing that XRN2 depletion leads to increased R-loop accumulation and that XRN2 is enriched at R-loop-prone genomic regions^31,78–80^.

RNA decapping prior to XRN2-mediated degradation raises intriguing questions: What decapping enzyme(s) operate at DSBs? Are canonical cytoplasmic decapping factors, such as DCP2, recruited to nuclear damage sites by LSm4 condensates, or does a specialized nuclear decapping machinery exist? Supporting the former possibility, a recent study demonstrated that XRN2 immunoprecipitated from HeLa nuclear extracts co-purifies with DCP2, DCP1, and EDC3, suggesting the existence of a nuclear decapping complex associated with XRN2 ^81^. Alternatively, DXO enzyme, which selectively decaps nascent transcripts bearing incomplete 5’ terminal caps in the nucleus and possesses 5’→3’ exoribonuclease activity, represents another candidate for a decapping enzyme at DSBs ^82^. Our findings position LSm4 as a potential scaffold that coordinates nuclear decapping activities at DSBs. Future studies are required to elucidate the mechanism underlying RNA decapping at DNA damage sites.

Beyond the LSm4-XRN2 axis operating in the 5’→3’ direction, the nuclear exosome subunit EXOSC10 is recruited to DSBs to degrade damage-induced lncRNAs from the 3’ end, facilitating R-loop resolution and DNA end resection ^83^. Whether LSm4 condensates are prerequisite for EXOSC10 accumulation at damage sites or whether these represent independent parallel pathways for RNA clearance at DSBs remains to be determined.

The LSm4-XRN2 axis complements the known mechanisms of R-loop resolution at DSBs. Helicases such as SETX actively unwind R-loops, liberating the RNA strand from DNA. However, if this displaced RNA persists in the vicinity of the break, it can rapidly re-anneal to DNA, reforming deleterious R-loops. Our findings indicate that LSm4-driven degradation of nascent RNA provides a surveillance mechanism; by rapidly clearing the freed RNA strand, the cell prevents R-loop re-accumulation and ensures that the single-stranded DNA necessary for RAD51 loading remains accessible. This model is strongly supported by our observation that targeted resolution of R-loops at a specific DSB using dCas9-RNaseH1 rescues RAD51 recruitment in LSm4-deficient cells. These data demonstrate that defective RAD51 loading in LSm4-deficient cells results from excessive accumulation of R-loops at DSBs.

While our study focused on LSm4 function at DSBs, other transcription-associated DNA lesions caused by UV or alkylating agents also generate DNA-RNA conflicts that require resolution. This suggests that RNA clearance may be a general prerequisite for repairing various types of DNA lesions at transcriptionally active regions. Whether LSm4 condensates act at these various lesion types remains to be determined.

Interestingly, emerging evidence implicates LSm4 dysregulation in tumorigenesis. LSm4 is significantly overexpressed in hepatocellular carcinoma, breast cancer, non-small cell lung cancer, ovarian cancer, and pancreatic cancer, where high expression correlates with advanced clinical stage, lymph node metastasis, and poor overall survival. How LSm4 dysregulation affects cancer progression remains poorly understood. Based on our findings, we speculate that LSm4 dysregulation creates a permissive environment for genomic instability by disrupting the balance between RNA metabolism and DNA repair. Given this emerging connection between LSm4-mediated RNA decay at DSBs and its frequent alterations in cancer, LSm4 may represent both a prognostic biomarker and a potential therapeutic target, particularly in HR-deficient tumors ^84–91^.

## Data availability

All ChIP-seq data generated in this study have been deposited at ArrayExpress: E-MTAB- 16861, and all RNA-seq data generated in this study have been deposited at ArrayExpress: E-MTAB-16857. Accession numbers are listed in the Key Resources Table. This paper does not report original code. Any additional information required to reanalyze the data reported in this paper is available from the lead contact upon request.

## Acknowledgements

We thank Nitzan Dahan, Yael Lupu Haber, Tally Chalolachvilli-Mergener, Aviv Lutaty and Yousef Mansour from the Life Sciences and Engineering Infrastructure Unit at the Technion for their assistance with microscopy and flow cytometry. ChIP-seq and RNA-seq library preparation and sequencing were performed Azrieli Technion Genomics Center, Technion-Israel Institute of Technology, Haifa, Israel. We acknowledge the support of the Azrieli Foundation for funding this research. We are grateful to members of the Ayoub lab for critical reading of the manuscript. We thank BioRender (biorender.com) for assistance in preparing the graphical model and figure schematics. Research in the Ayoub lab is supported by grants from the Israel Science Foundation (2511_2019 and 1962_2024) and Israel Cancer Research Fund (22-110-PG). We acknowledge the support of the Azrieli Foundation for funding this research. M.M.D. is supported by the Neubauer Family fellowship. C.E. is supported by VATAT scholarship program for outstanding master students. A.S.B is supported by the RTICC fellowship for multidisciplinary cancer research. N.A. is supported by the Neubauer Family foundation. N.A. also acknowledges the support of Chil and Berta Weissman Chair in Precision Medicine.

## Author Contributions

M.M.D performed and analyzed all the experiments in this work except of the experiments described below. Also, wrote the materials and methods and helped in writing the manuscript. E.R.A help in conceiving and planning the described experiments, performed the experiments described in Figures 1G and 5C. L.A.B performed the experiments described in Figures 1A, 1D, 2F-J, 3G-K, 4, S2 and 5A. A.S.B performed the conservation analysis described in Figures 1I, 1J and analyzed all RNA-seq data described in Figures 4B-F and Supplementary Figure 2. F.E.M. analyzed the LSm4 ChIP-seq data described in Figure 2C. C.E. performed the experiment described in Figure S1E. N.A. conceived the study, planned the experiments, analyzed the data and wrote the original draft.

## STAR Methods

### Key resources table

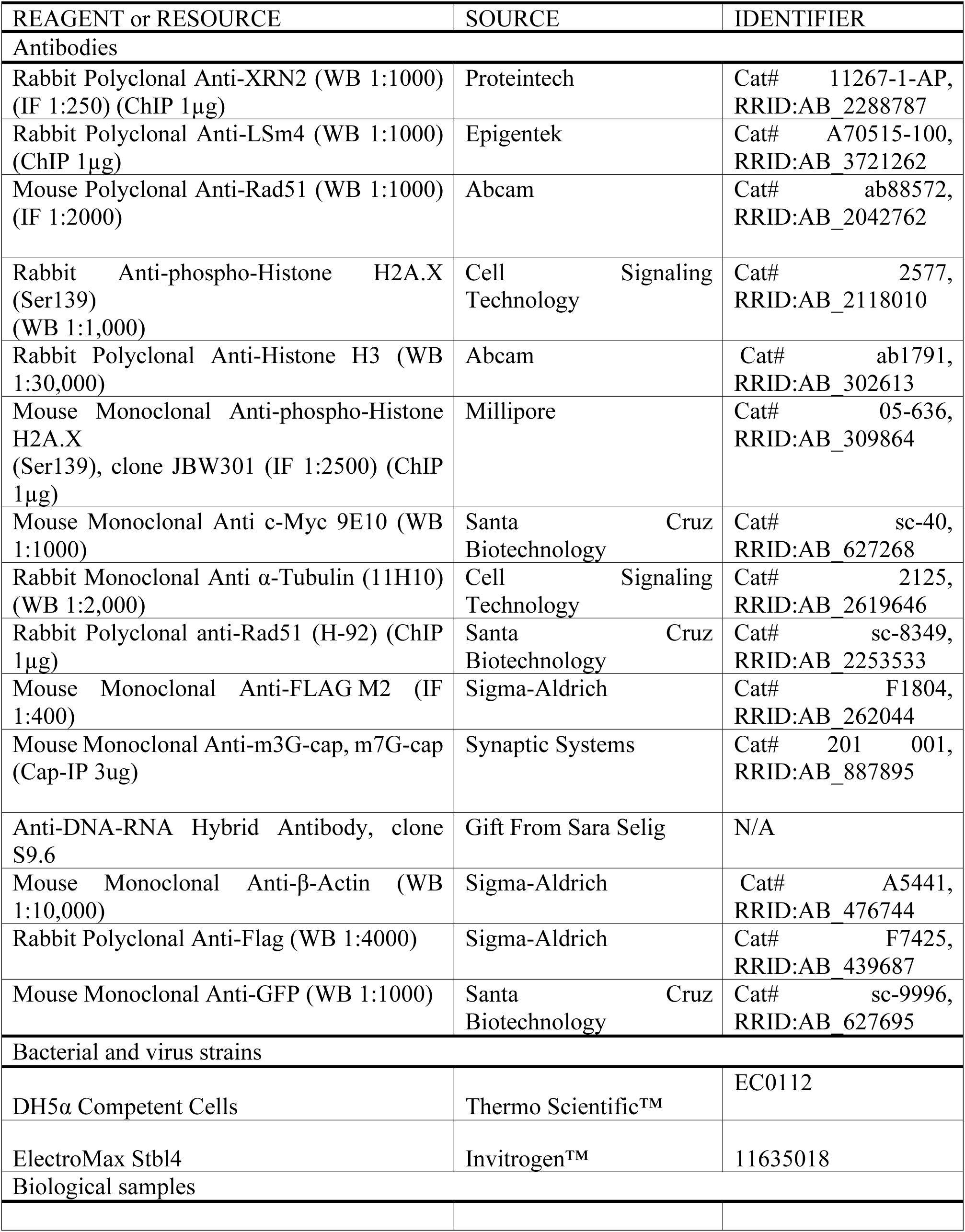
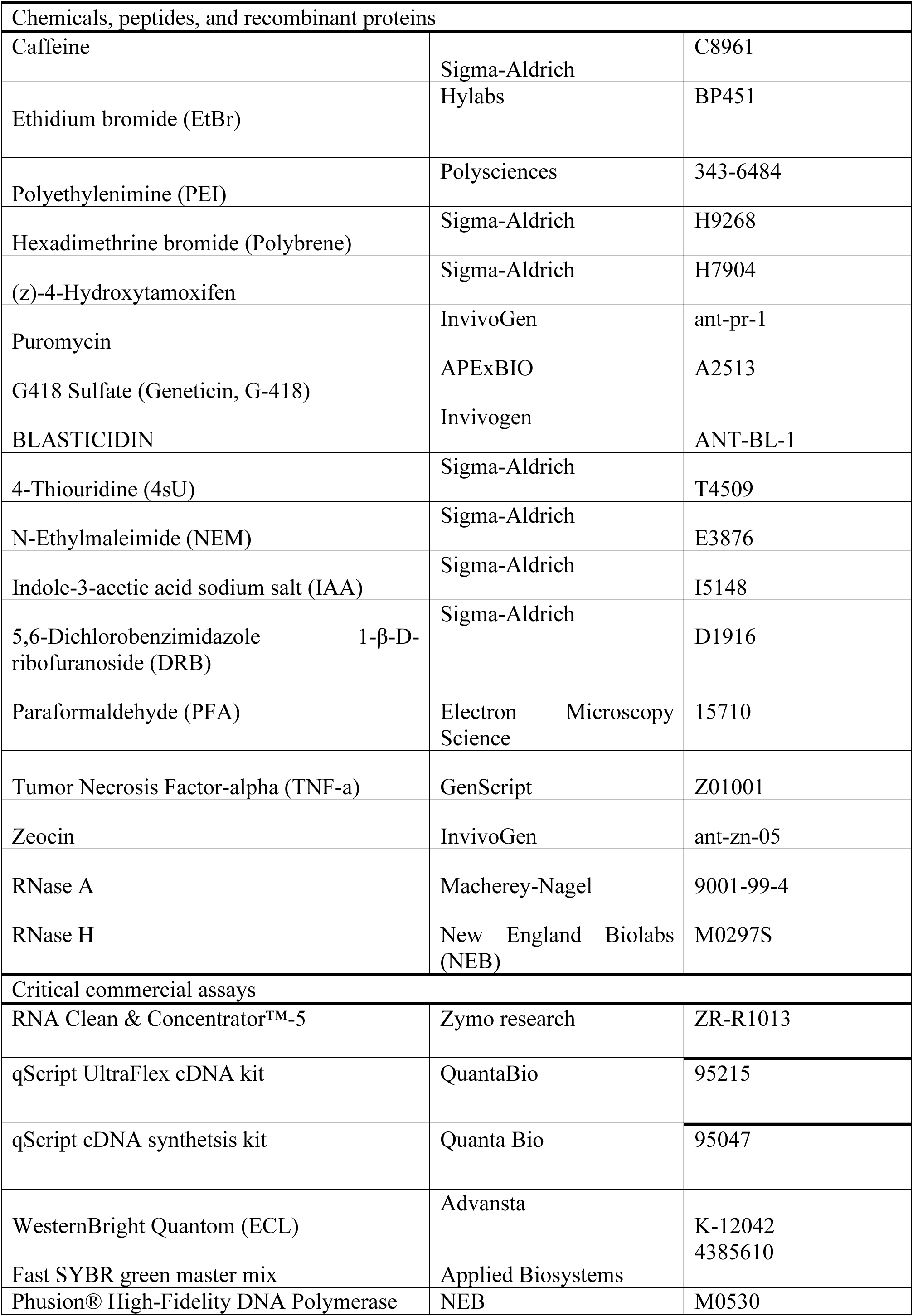
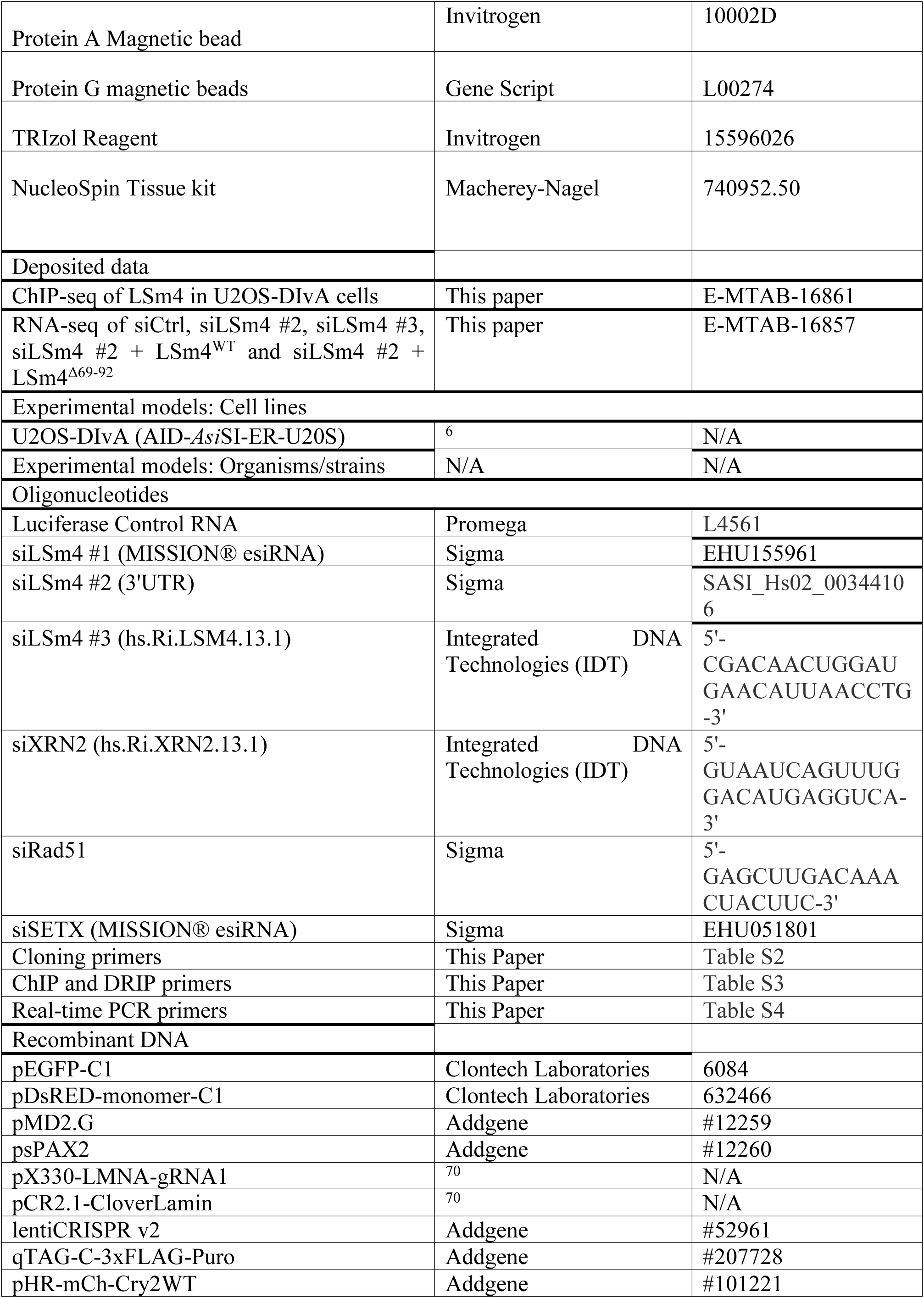
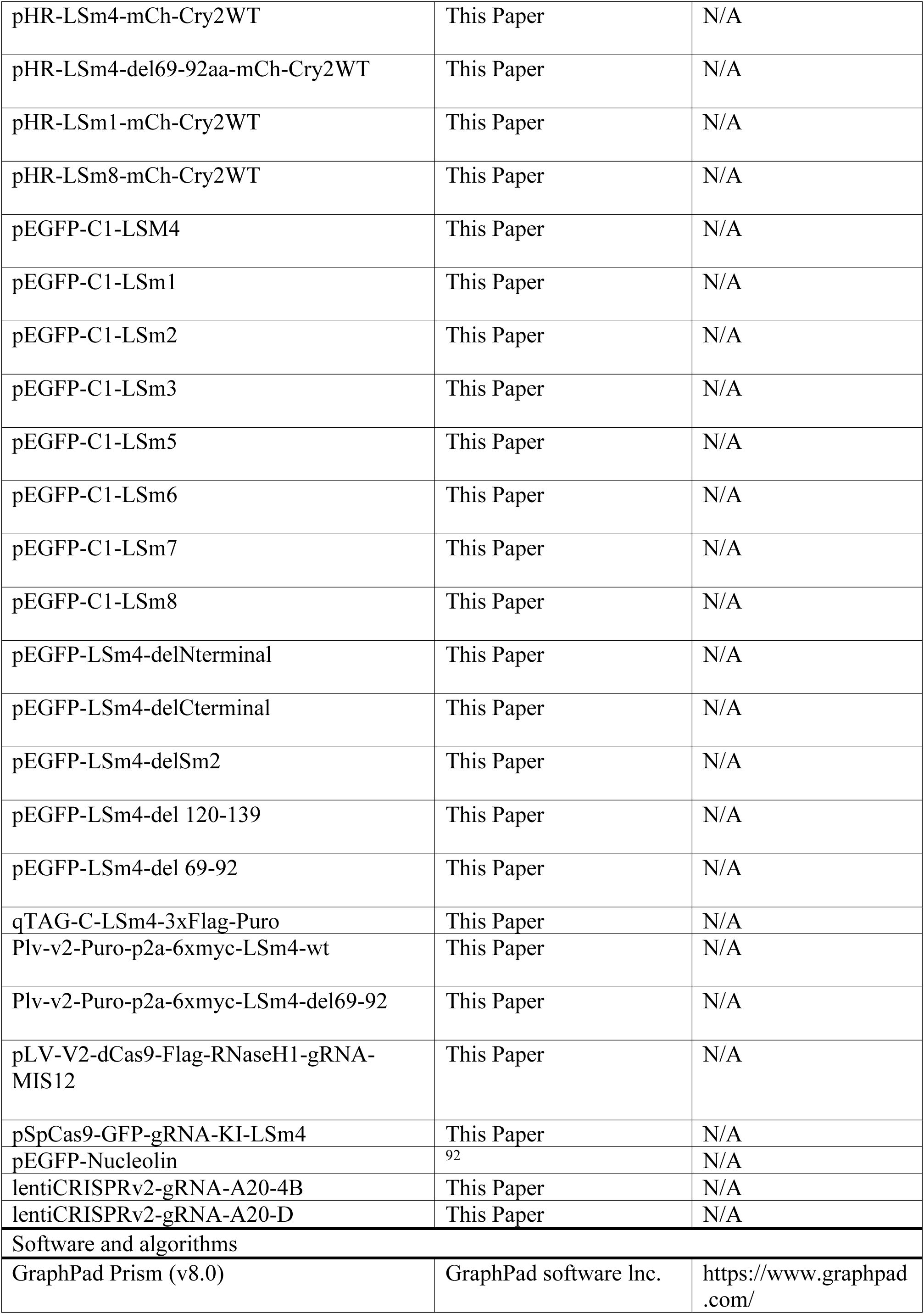
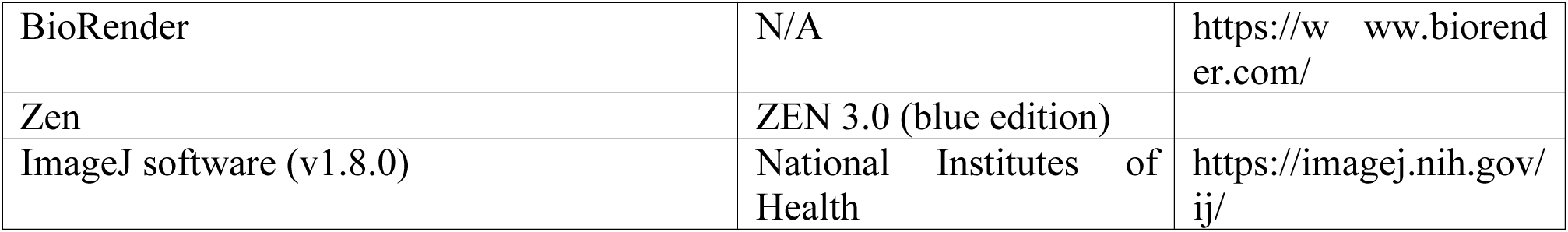

### Experimental model and subject details

U2OS, HEK293T and HeLa cell lines were grown in Dulbecco’s Modified Eagle’s Medium (DMEM; Gibco) supplemented with 10% heat-inactivated fetal bovine serum (FBS), 2 mM L-glutamine, 100 units/mL penicillin, and 100 μg/mL streptomycin (Gibco). U2OS-DIvA cell line (expressing AID-*Asi*SI-ER) was cultured in DMEM supplemented with the standard components described above, together with 1 mM sodium pyruvate and 400 μg/mL G418 for selection. U2OS-DIvA cells stably transduced with lentiviral vectors expressing LSm4^WT^ or LSm4^Δ69–92^ were maintained under 0.6 μg/mL puromycin selection.

### Cell culture and transfections

For *Asi*SI-dependent DSB induction, cells were treated with 300 nM (Z)-4-hydroxytamoxifen (4-OHT) for 4 hours. Plasmid DNA transfections were performed using polyethylenimine (PEI) following the manufacturer’s instructions. siRNA transfections were carried out using Lipofectamine RNAiMax (Thermo Fisher Scientific) according to the manufacturer’s protocol. The siRNA sequences used in this study are listed in the Key Resources Table.

### Generation of lentiviral particles and cell transduction

For rescue experiments, LSm4^WT^ and LSm4^Δ69-92^ were cloned into the lentiCRISPR-v2 lentiviral vector (Key Resource Table). To produce viral particles, HEK293T cells seeded in 10 cm dishes were co-transfected with 1.64 pmol of the lentiviral construct together with the packaging plasmids psPAX2 (1.3 pmol) and pMD2.G (0.72 pmol). Viral supernatants were collected at 48 and 72 hours post-transfection, pooled, and passed through 0.45 μm filters. The resulting viral particles were used to transduce U2OS-DIvA or U2OS cells, which were subsequently maintained under 0.6 μg/mL puromycin selection.

### OptoDroplet assay

OptoDroplet assays were performed as described ^58^. LSm1, LSm8, LSm4^WT^, and LSm4^Δ69–92^ were individually cloned into the pHR-mCherry-Cry2WT vector (Key Resource Table). Viral particles were generated in HEK293T cells as described above. U2OS cells transduced with pHR-mCherry-Cry2 constructs were seeded on FluoroDishes (Ibidi; Cat# 81158). At 48 hours post-transduction, cells were subjected to light activation and live imaging using an LSM-700 inverted confocal microscope equipped with a 37°C/5% CO2 environmental chamber. mCherry-positive cells were stimulated with 488 nm light (10% laser power, 15 iterations) to drive Cry2 oligomerization. Time-lapse images of mCherry fluorescence were acquired every 10 seconds using 555 nm excitation (6% laser power).

### Western blot

Total protein extracts were prepared using hot-lysis buffer (50 mM Tris-HCl pH 7.5, 5 mM EDTA, 1% SDS) supplemented with protease inhibitor cocktail (Calbiochem), as previously described ^93^. Proteins were resolved by SDS-PAGE, transferred to nitrocellulose membranes, and probed with the antibodies indicated in each figure at the dilutions listed in the Key Resources Table. H3, β-actin, and tubulin served as loading controls; γH2AX was used as a marker of DNA damage induction. Band intensities were quantified using ImageJ software and normalized to the respective loading control. Protein molecular weight sizes are indicated on the right side of each blot.

### Immunofluorescence

Cells were grown on coverslips for 24 hours and processed for immunofluorescence as previously described ^93^. Cells were immunostained with the antibodies indicated in each figure (see Key Resources Table for dilutions). Slides were imaged on an inverted Zeiss LSM-700 confocal microscope equipped with a 40× oil-immersion objective. For detection of R-loops using GFP-dRNaseH1 ^69^, cells were fixed with ice-cold methanol for 5 min at −20°C, followed by permeabilization with 0.25% Triton X-100 in PBS. Samples were blocked in staining buffer (3% BSA in PBS) for 30 min, then incubated with purified GFP-dRNaseH1 (0.188 μg/mL; 1:2,000 dilution) for 1.5 hours at 37°C. After washing, coverslips were mounted with DAPI-containing mounting medium.

### RNA isolation, reverse transcription, and quantitative real-time PCR

Total RNA was isolated using TRIzol reagent (Ambion) according to the manufacturer’s instructions. 1 μg of RNA was reverse transcribed using the qScript cDNA Synthesis Kit (Quanta Bio) with random primers. Quantitative real-time PCR was performed on a StepOnePlus Real-Time PCR System (Applied Biosystems) using Fast SYBR Green Master Mix (Applied Biosystems) and the primers listed in Table S4. Relative expression was calculated by the ΔΔCT method. GAPDH or 18S rRNA was used as a reference gene as indicated in the figure legends.

### RNA sequencing (RNA-seq) and analysis

Three biological replicates of RNA samples were purified from U2OS cells treated as described. Total RNA was isolated using TRIzol reagent (Ambion) according to the manufacturer’s instructions. RNA sequencing libraries were prepared using NEBNext Ultra II Directional RNA Library Prep Kit for Illumina (NEB, cat no. E7760) . Sequencing was performed at the Azrieli Technion Genomics Center, Technion – Israel Institute of Technology using Illumina NextSeq2000, using P3 XLEAP-SBS 300 cycles (Read1-150; Index1-8; Index2-8; Read2-150) (Illumina, cat no. 20100988) sequencing kit . Average read depth was 80 million reads per sample. For differential gene expression analysis, raw sequencing reads were aligned to GENCODE GRCh38 genome assembly using the splice-sensitive aligner HISAT2^94^ and differential gene analysis was performed in R using the DESeq2 package^95^. Pathway enrichment analysis and gene ontology was conducted using ShinyGO^96^. To analyze alternative splicing events, HISAT2-aligned reads were subjected to alternative splicing events detection using rMATS package^97^.

### Tagging endogenous LSm4 using CRISPR-Cas9

CRISPR-Cas9-mediated tagging was used to introduce a 3×Flag tag at the C-terminus of the endogenous *LSm4* locus. U2OS cells were co-transfected with the pSpCas9-GFP-gRNA-KI- LSm4 plasmid (encoding SpCas9 and a gRNA targeting the *LSm4* coding sequence immediately upstream of the stop codon) and a donor plasmid qTAG-C-LSm4-3xFlag-Puro encoding left and right homology arms flanking the 3×Flag sequence, a P2A self-cleavage peptide, and a puromycin resistance cassette (Key Resource Table). Puromycin-resistant clones were screened by genomic PCR using the diagnostic primers listed in Key Resource Table.

### Laser microirradiation

Laser microirradiation was performed as previously described ^93^. Briefly, U2OS cells were seeded on FluoroDishes (Ibidi; Cat# 81158) and sensitized with Hoechst 33342 (1 μg/mL) for 10 min at 37°C. Microirradiation was carried out using an LSM-700 inverted confocal microscope equipped with a 37°C/5% CO2 environmental chamber. DNA damage was induced by targeting a single nuclear region with a 405 nm laser (20 iterations). Live-cell imaging was performed with 488 nm excitation, and fluorescence intensity at irradiated sites was quantified using ZEN software (Zeiss).

### Fluorescence recovery after photobleaching (FRAP)

U2OS cells expressing EGFP-LSm4 were seeded on FluoroDishes (Ibidi; Cat# 81158) and cultured for 12–16 hours prior to imaging. FRAP experiments were performed on an inverted Zeiss LSM-700 confocal microscope with a 37°C/5% CO2 environmental chamber. A single nucleolus per cell was photobleached using a 488 nm laser at 100% power (40 iterations). Fluorescence recovery was monitored by time-lapse imaging at the indicated intervals. Intensities within the bleached region were quantified using ZEN software and normalized by setting the pre-bleach value to 100%.

### Sequence conservation analysis

Human LSm4 protein sequence was obtained from UniProt. Homologous sequences from selected species were retrieved and aligned using MAFFT v7 with the L-INS-i iterative refinement algorithm and default BLOSUM62 scoring parameters ^98,99^.

### A20 transcriptional activation system

HeLa cells were left untreated or treated with TNF-α (10 ng/mL) for 48 hours to activate *A20* transcription. Cells were subsequently transfected with expression vectors encoding SpCas9 and site-specific gRNAs to introduce DSBs at defined positions within the *A20* locus. At 48 hours post-transfection, cells were harvested for chromatin immunoprecipitation (ChIP) using anti-LSm4 antibody, with anti-γH2AX serving as a positive control for DSB induction. The lentiCRISPRv2-gRNA-A20-4B and lentiCRISPRv2-gRNA-A20-D plasmids were used to introduce DSB-1 and DSB-2, respectively (Key Resource Table). ChIP primer sequences are listed in Table S3.

### Transcriptional shutoff assay

To measure RNA stability at DSB-proximal loci, DIvA cells transfected with siLSm4 or siCtrl (48 hours) were treated with 300 nM 4-OHT for 4 hours to induce DSBs. Transcription was then globally inhibited by addition of 100 μM 5,6-dichlorobenzimidazole 1-β-D-ribofuranoside (DRB; Sigma-Aldrich, D1916) for the indicated time points. Total RNA was extracted using TRIzol, and 1 μg was reverse transcribed using the qScript cDNA Synthesis Kit (Quanta Bio) with random hexamer primers. mRNA levels of DSB-proximal transcripts at the *MIS12* and *AGA* loci were quantified by RT-qPCR using primers listed in Key Resource Table and normalized to 18S rRNA. Relative expression was calculated using the ΔΔCT method.

### Roadblock-qPCR

Roadblock-qPCR was performed as previously described, with minor adaptations ^67^. DIvA cells transfected with siLSm4 or siCtrl (48 hours) were treated with 300 nM 4-OHT for 4 hours to induce DSBs. Newly synthesized RNA was pulse-labeled by incubating cells with 400 μM 4-thiouridine (4sU; Sigma-Aldrich, T4509) for the indicated times. Total RNA was extracted, and 1 μg was spiked with a fixed amount of luciferase mRNA (Promega, L4561) as an exogenous normalization control. RNA was treated with N-ethylmaleimide (NEM; Sigma-Aldrich, E3876) to selectively block reverse transcription of 4sU-labeled transcripts and then purified using the RNA Clean & Concentrator-5 kit (Zymo Research, ZR-R1013). First-strand cDNA synthesis was performed with the qScript UltraFlex cDNA Kit (QuantaBio, 95215) using oligo(dT) primers. RT-qPCR was conducted on a StepOnePlus system (Applied Biosystems) using Fast SYBR Green Master Mix and primers targeting RBMXL1 and MIS12 transcripts (Key Resource Table). RNA abundance was normalized to either 18S rRNA or the luciferase spike-in control.

### DNA:RNA immunoprecipitation followed by quantitative PCR (DRIP-qPCR)

DRIP-qPCR was performed as previously described (Sanz and Chédin, 2019), with minor modifications. DIvA cells transfected with siLSm4 or siCtrl (48 hours) were treated with 300 nM 4-OHT for 4 hours. Cells were trypsinized, pelleted, and washed once with PBS. Nucleic acids were extracted overnight at 37°C in TE buffer (pH 8.0) containing 0.5% SDS and proteinase K (20 mg/mL) and purified by phenol/chloroform/isoamyl alcohol (25:24:1) extraction and ethanol precipitation. DNA was digested overnight at 37°C with a restriction enzyme cocktail (EcoRI, HindIII, BsrGI, XbaI, SspI; NEB), with or without RNase H treatment, followed by a second phenol/chloroform extraction and ethanol precipitation. From 4.4 μg of digested DNA diluted in 500 μL TE, 50 μL was reserved as input. The remaining 450 μL was supplemented with 50 μL 10× binding buffer (100 mM NaPO4 pH 7.0, 1.4 M NaCl, 0.5% Triton X-100) and 20 μL S9.6 antibody, then incubated overnight at 4°C with rotation. Protein G magnetic beads (GeneScript; L00274) pre-washed in 1× binding buffer were added and samples were incubated for 3 hours at 4°C. Beads were washed three times with 1× binding buffer (10 min each at room temperature), resuspended in elution buffer with proteinase K (20 mg/mL), and incubated at 55°C for 45 min. DRIP DNA was purified by phenol/chloroform extraction and ethanol precipitation, then resuspended in RNase-free TE buffer. qPCR was performed using primers for *SNRNP* (R-loop-negative control), *RPL13A* (R-loop-positive control), and *Asi*SI-proximal loci DSB-I (*RBMXL1*), DSB-II (*MIS12*), and DSB-IV (*AGA*) (Key Resource Table).

### Dot blot

Genomic DNA was extracted from DIvA cells transfected with siCtrl or siLSm4 as described for DRIP-qPCR. Purified, digested DNA was serially diluted (2-fold, starting from 2 μg) and spotted onto positively charged nylon membranes (Amersham Hybond-N+; RPN203B). Membranes were UV cross-linked (1,200 μJ/cm2), blocked in 5% milk in TBST for 1 hour at room temperature, and incubated overnight at 4°C with S9.6 antibody. After washing with TBST, the membrane was incubated with secondary antibody for 45 min at room temperature, and signal was detected using the Quantum ECL kit. Methylene blue staining served as a loading control. RNase H1-pretreated samples were used as specificity controls.

### Chromatin immunoprecipitation followed by quantitative PCR (ChIP-qPCR)

ChIP assays were carried out as previously described ^7^. DIvA cells plated in 150 mm dishes were treated with 300 nM 4-OHT for 4 hours upon reaching ∼80% confluency. Cells were crosslinked with 1% paraformaldehyde (PFA) for 10 min at room temperature, and crosslinking was quenched with 0.125 M glycine for 5 min. Following lysis, chromatin was sheared to fragments of 300–500 bp using a Vibra-Cell sonicator (15 sec ON / 30 sec OFF, 35% duty cycle, 20 cycles). Five percent of each supernatant was reserved as input. Chromatin was immunoprecipitated with 2 μg of the indicated antibodies and Protein A/G magnetic beads. After reverse crosslinking, precipitated DNA was purified using the Macherey-Nagel NucleoSpin kit. Both input and IP samples were analyzed by qPCR using the primers listed in Key Resource Table. ChIP enrichment is expressed as percent of input and is presented as mean ± SD from three independent experiments.

### Isolation of chromatin-associated RNA

Cells were trypsinized and washed once with cold PBS. Cytoplasmic RNA was first separated using Buffer A (10 mM HEPES pH 7.5, 1.5 mM MgCl2, 10 mM KCl, 0.5 mM DTT, 1 mM PMSF, protease inhibitors, 0.25% NP-40) for 7 min on ice, followed by centrifugation at 2,000 × g for 3 min. The nuclear soluble fraction was then extracted by incubating the nuclear pellet in Buffer C (20 mM HEPES pH 7.5, 10% glycerol, 0.42 M KCl, 4 mM MgCl2, 0.2 mM EDTA, 0.5 mM DTT, 1 mM PMSF, protease inhibitors) for 30 min on ice. Chromatin was pelleted by centrifugation at maximum speed at 4°C, and chromatin-associated RNA was extracted from the pellet using TRIzol reagent according to the manufacturer’s instructions.

### Cap immunoprecipitation (Cap-IP)

Cap-IP was performed as previously described ^100^, with modifications. DIvA cells were transfected with siLSm4 or siCtrl for 48 hours, followed by 4-OHT treatment for 4 hours. Chromatin-associated RNA was isolated as described above. Protein G magnetic beads were washed three times with PBS and once with Cap-IP buffer (0.05% BSA, 1 mM DTT, 0.05% Triton X-100), then pre-loaded with 3 μg of anti-m7G antibody (Synaptic Systems, #201-001) in 200 μL Cap-IP buffer for 3 hours at room temperature. 10 μg of chromatin-associated RNA was heat-denatured at 65°C for 5 min, snap-cooled on ice for 5 min, and added to the antibody-bead complex for 1 hour at room temperature. After four washes with Cap-IP buffer, capped RNA was eluted and purified using the RNA Clean & Concentrator-5 kit (Zymo Research, ZR- R1013). Capped RNA levels were quantified by RT-qPCR and normalized to input.

### ChIP-seq analysis

ChIP-seq samples were aligned to the GRCh38/hg38 human genome assembly using Bowtie2 ^101^. Aligned reads were converted to BAM format and sorted, indexed, and quality-filtered using SAMtools ^102^. Coverage tracks were generated using bamCoverage (deepTools) ^103^ with RPKM normalization. Coverage bigWig files from biological replicates were merged using bigwigCompare with the ’add’ operation. Average log2 ratios between −4-OHT and +4-OHT conditions were computed using bigwigCompare with the ’log2’ operation. Averaged ChIP-seq log2 profiles around *Asi*SI sites in transcriptionally active or silent/intergenic regions (as defined in ^34^) were computed using computeMatrix (deepTools) with 500 bp smoothing bins in a 5 kb window. Data are presented as dot plots in which each point represents the average log2 (−4-OHT/+4-OHT) value at a single *Asi*SI site. Statistical significance was assessed by nonparametric unpaired Mann-Whitney-Wilcoxon test.

### Biochemical fractionation

Biochemical fractionation was performed as previously described ^23^, with minor modifications. U2OS or DIvA cells were left untreated or exposed to DNA-damaging agents as indicated in figure legends. Cells were lysed in cytoplasmic lysis buffer (Buffer A) for 5 min at 4°C, and lysates were centrifuged at 1,500 × g for 5 min at 4°C to pellet nuclei; the supernatant constituted the cytoplasmic fraction. The nuclear pellet was resuspended in nuclear lysis buffer (Buffer B: 3 mM EDTA, 0.2 mM EGTA, 1 mM DTT, 1 mM PMSF, and protease inhibitor cocktail) and incubated for 10 min on ice, followed by centrifugation at 1,700 × g for 5 min at 4°C to collect the nuclear soluble fraction. The chromatin-bound fraction was recovered by resuspending the remaining pellet in hot-lysis buffer (1% SDS, 5 mM EDTA, 50 mM Tris-HCl pH 7.5, protease inhibitors), boiling for 15 min, sonicating with two 15-second pulses at 35% amplitude, and centrifuging at 16,000 × g for 20 min at 12°C. All fractions were analyzed by western blot.

### Endogenous homologous recombination assay (mClover)

HR efficiency was measured using a Cas9-mediated mClover knock-in assay, as previously described ^70^. Briefly, cells were seeded in 6-well plates and co-transfected with 1.6 μg of pX330-LMNA-gRNA1 (encoding Cas9 and a gRNA targeting exon 1 of *LMNA*) and 0.4 μg of pCR2.1-CloverLamin (HR donor template). 0.4 μg of pDsRed-Monomer-C1 was included as a transfection efficiency control. Where indicated, caffeine (4 mM) was added 16 hours post-transfection as a negative control for HR. At 48 hours post-transfection, cells were harvested and analyzed by flow cytometry on a Cytek Aurora instrument. HR efficiency was calculated as the percentage of mClover-positive cells among DsRed-Monomer-positive (transfected) cells. Data are presented as mean ± SD from at least three independent experiments.

### Translocation assay

Chromosomal translocation frequency was assessed as previously described ^43^. AID-DIvA cells transfected with siLSm4 or siCtrl (48 hours) were treated with 300 nM 4-OHT for 4 hours to induce DSBs. Cells were then washed three times with PBS and treated with 500 μg/mL indole-3-acetic acid (IAA) for 2 hours to degrade AID-*Asi*SI and allow DSB repair to proceed. Illegitimate rejoining events between DSBs at the *MIS12* (chr17:5,390,209) and *LYRM2* (chr6:9,034,817) loci were quantified by qPCR using the primers listed in Key Resource Table. qPCR signals were normalized to a reference amplicon (*Norm17*) located distal to *Asi*SI cleavage sites and γH2AX-enriched domains. Relative translocation frequencies were calculated using the ΔΔCt method.

### Quantification and statistical analysis

Statistical analyses were performed using GraphPad Prism 8.0. Data are presented as mean ± SD unless otherwise stated. Comparisons between two groups were carried out using two-tailed unpaired Student’s t-tests. For ChIP-seq data, nonparametric unpaired Mann-Whitney-Wilcoxon tests were used to assess differences between *Asi*SI site categories. Statistical significance is indicated as follows: *P < 0.05; **P < 0.01; ***P < 0.001; ****P < 0.0001; ns, not significant. The number of independent experiments is indicated in each figure legend.

## Declaration of generative AI and AI-assisted technologies in the writing process

During the preparation of this work the authors used ChatGPT for grammatical proofreading and stylistic improvement. After using this tool, the authors reviewed and edited the content as needed and take full responsibility for the content of the published article.

## References

1. Heyer, W.D., Ehmsen, K.T., and Liu, J. (2010). Regulation of homologous recombination in eukaryotes. Annual review of genetics 44, 113–139. 10.1146/annurev-genet-051710-150955.

2. Ceccaldi, R., and Cejka, P. (2025). Mechanisms and regulation of DNA end resection in the maintenance of genome stability. Nat Rev Mol Cell Biol. 10.1038/s41580-025-00841-4.

3. Ceccaldi, R., Rondinelli, B., and D’Andrea, A.D. (2016). Repair Pathway Choices and Consequences at the Double-Strand Break. Trends Cell Biol 26, 52-64. 10.1016/j.tcb.2015.07.009.

4. Scully, R., Panday, A., Elango, R., and Willis, N.A. (2019). DNA double-strand break repair-pathway choice in somatic mammalian cells. Nat Rev Mol Cell Biol 20, 698–714. 10.1038/s41580-019-0152-0.

5. Marnef, A., Cohen, S., and Legube, G. (2017). Transcription-Coupled DNA Double-Strand Break Repair: Active Genes Need Special Care. J Mol Biol 429, 1277-1288. 10.1016/j.jmb.2017.03.024.

6. Iacovoni, J.S., Caron, P., Lassadi, I., Nicolas, E., Massip, L., Trouche, D., and Legube, G. (2010). High-resolution profiling of gammaH2AX around DNA double strand breaks in the mammalian genome. EMBO J 29, 1446-1457. 10.1038/emboj.2010.38.

7. Aymard, F., Bugler, B., Schmidt, C.K., Guillou, E., Caron, P., Briois, S., Iacovoni, J.S., Daburon, V., Miller, K.M., Jackson, S.P., and Legube, G. (2014). Transcriptionally active chromatin recruits homologous recombination at DNA double-strand breaks. Nat Struct Mol Biol 21, 366–374. 10.1038/nsmb.2796.

8. Ouyang, J., Lan, L., and Zou, L. (2017). Regulation of DNA break repair by transcription and RNA. Sci China Life Sci 60, 1081–1086. 10.1007/s11427-017-9164-1.

9. Tang, J., Cho, N.W., Cui, G., Manion, E.M., Shanbhag, N.M., Botuyan, M.V., Mer, G., and Greenberg, R.A. (2013). Acetylation limits 53BP1 association with damaged chromatin to promote homologous recombination. Nat Struct Mol Biol 20, 317–325. 10.1038/nsmb.2499.

10. Wei, L., Nakajima, S., Bohm, S., Bernstein, K.A., Shen, Z., Tsang, M., Levine, A.S., and Lan, L. (2015). DNA damage during the G0/G1 phase triggers RNA-templated, Cockayne syndrome B-dependent homologous recombination. Proc Natl Acad Sci U S A 112, E3495–3504. 10.1073/pnas.1507105112.

11. Teng, Y., Yadav, T., Duan, M., Tan, J., Xiang, Y., Gao, B., Xu, J., Liang, Z., Liu, Y., Nakajima, S., et al. (2018). ROS-induced R loops trigger a transcription-coupled but BRCA1/2-independent homologous recombination pathway through CSB. Nature communications 9, 4115. 10.1038/s41467-018-06586-3.

12. Chen, H., Yang, H., Zhu, X., Yadav, T., Ouyang, J., Truesdell, S.S., Tan, J., Wang, Y., Duan, M., Wei, L., et al. (2020). m(5)C modification of mRNA serves a DNA damage code to promote homologous recombination. Nature communications 11, 2834. 10.1038/s41467-020-16722-7.

13. Ouyang, J., Yadav, T., Zhang, J.M., Yang, H., Rheinbay, E., Guo, H., Haber, D.A., Lan, L., and Zou, L. (2021). RNA transcripts stimulate homologous recombination by forming DR-loops. Nature 594, 283-288. 10.1038/s41586-021-03538-8.

14. Palancade, B., and Rothstein, R. (2021). The Ultimate (Mis)match: When DNA Meets RNA. Cells 10. 10.3390/cells10061433.

15. Ortega, P., Merida-Cerro, J.A., Rondon, A.G., Gomez-Gonzalez, B., and Aguilera, A. (2021). DNA-RNA hybrids at DSBs interfere with repair by homologous recombination. eLife 10. 10.7554/eLife.69881.

16. Stoimenov, I., Schultz, N., Gottipati, P., and Helleday, T. (2011). Transcription inhibition by DRB potentiates recombinational repair of UV lesions in mammalian cells. PLoS One 6, e19492. 10.1371/journal.pone.0019492.

17. Marini, F., Rawal, C.C., Liberi, G., and Pellicioli, A. (2019). Regulation of DNA Double Strand Breaks Processing: Focus on Barriers. Front Mol Biosci 6, 55. 10.3389/fmolb.2019.00055.

18. Belotserkovskii, B.P., Tornaletti, S., D’Souza, A.D., and Hanawalt, P.C. (2018). R-loop generation during transcription: Formation, processing and cellular outcomes. DNA Repair (Amst) 71, 69–81. 10.1016/j.dnarep.2018.08.009.

19. Nudler, E. (2012). RNA polymerase backtracking in gene regulation and genome instability. Cell 149, 1438-1445. 10.1016/j.cell.2012.06.003.

20. Abu-Zhayia, E.R., Bishara, L.A., Machour, F.E., Barisaac, A.S., Ben-Oz, B.M., and Ayoub, N. (2022). CDYL1-dependent decrease in lysine crotonylation at DNA double-strand break sites functionally uncouples transcriptional silencing and repair. Mol Cell 82, 1940–1955 e1947. 10.1016/j.molcel.2022.03.031.

21. Machour, F.E., and Ayoub, N. (2020). Transcriptional Regulation at DSBs: Mechanisms and Consequences. Trends Genet 36, 981-997. 10.1016/j.tig.2020.01.001.

22. Abu-Zhayia, E.R., Machour, F.E., and Ayoub, N. (2019). HDAC-dependent decrease in histone crotonylation during DNA damage. J Mol Cell Biol 11, 804–806. 10.1093/jmcb/mjz019.

23. Abu-Zhayia, E.R., Awwad, S.W., Ben-Oz, B.M., Khoury-Haddad, H., and Ayoub, N. (2018). CDYL1 fosters double-strand break-induced transcription silencing and promotes homology-directed repair. J Mol Cell Biol 10, 341–357. 10.1093/jmcb/mjx050.

24. Awwad, S.W., Abu-Zhayia, E.R., Guttmann-Raviv, N., and Ayoub, N. (2017). NELF-E is recruited to DNA double-strand break sites to promote transcriptional repression and repair. EMBO Rep 18, 745–764. 10.15252/embr.201643191.

25. Caron, P., van der Linden, J., and van Attikum, H. (2019). Bon voyage: A transcriptional journey around DNA breaks. DNA Repair (Amst) 82, 102686. 10.1016/j.dnarep.2019.102686.

26. Shanbhag, N.M., Rafalska-Metcalf, I.U., Balane-Bolivar, C., Janicki, S.M., and Greenberg, R.A. (2010). ATM-dependent chromatin changes silence transcription in cis to DNA double-strand breaks. Cell 141, 970–981.

27. Zatreanu, D., Han, Z., Mitter, R., Tumini, E., Williams, H., Gregersen, L., Dirac-Svejstrup, A.B., Roma, S., Stewart, A., Aguilera, A., and Svejstrup, J.Q. (2019). Elongation Factor TFIIS Prevents Transcription Stress and R-Loop Accumulation to Maintain Genome Stability. Mol Cell 76, 57-69 e59. 10.1016/j.molcel.2019.07.037.

28. Shivji, M.K.K., Renaudin, X., Williams, C.H., and Venkitaraman, A.R. (2018). BRCA2 Regulates Transcription Elongation by RNA Polymerase II to Prevent R-Loop Accumulation. Cell Rep 22, 1031–1039. 10.1016/j.celrep.2017.12.086.

29. De Santa, F., Totaro, M.G., Prosperini, E., Notarbartolo, S., Testa, G., and Natoli, G. (2007). The Histone H3 Lysine-27 Demethylase Jmjd3 Links Inflammation to Inhibition of Polycomb-Mediated Gene Silencing. Cell.

30. Crossley, M.P., Bocek, M., and Cimprich, K.A. (2019). R-Loops as Cellular Regulators and Genomic Threats. Mol Cell 73, 398–411. 10.1016/j.molcel.2019.01.024.

31. Skourti-Stathaki, K., Proudfoot, N.J., and Gromak, N. (2011). Human senataxin resolves RNA/DNA hybrids formed at transcriptional pause sites to promote Xrn2-dependent termination. Mol Cell 42, 794–805. 10.1016/j.molcel.2011.04.026.

32. Hatchi, E., Skourti-Stathaki, K., Ventz, S., Pinello, L., Yen, A., Kamieniarz-Gdula, K., Dimitrov, S., Pathania, S., McKinney, K.M., Eaton, M.L., et al. (2015). BRCA1 recruitment to transcriptional pause sites is required for R-loop-driven DNA damage repair. Mol Cell 57, 636–647. 10.1016/j.molcel.2015.01.011.

33. Marnef, A., and Legube, G. (2021). R-loops as Janus-faced modulators of DNA repair. Nat Cell Biol 23, 305–313. 10.1038/s41556-021-00663-4.

34. Saur, F., Lesage, E., Pradel, L., Collins, S., Finoux, A.L., Alghoul, E., Le Bozec, B., Rocher, V., Carette, R., Puget, N., et al. (2025). Transcriptional repression facilitates RNA:DNA hybrid accumulation at DNA double-strand breaks. Nat Cell Biol 27, 992–1005. 10.1038/s41556-025-01669-y.

35. Zhu, M., Wang, X., Zhao, H., and Wang, Z. (2025). Update on R-loops in genomic integrity: Formation, functions, and implications for human diseases. Genes Dis 12, 101401. 10.1016/j.gendis.2024.101401.

36. Garcia-Muse, T., and Aguilera, A. (2019). R Loops: From Physiological to Pathological Roles. Cell 179, 604-618. 10.1016/j.cell.2019.08.055.

37. Niehrs, C., and Luke, B. (2020). Regulatory R-loops as facilitators of gene expression and genome stability. Nat Rev Mol Cell Biol 21, 167–178. 10.1038/s41580-019-0206-3.

38. Alfano, L., Caporaso, A., Altieri, A., Dell’Aquila, M., Landi, C., Bini, L., Pentimalli, F., and Giordano, A. (2019). Depletion of the RNA binding protein HNRNPD impairs homologous recombination by inhibiting DNA-end resection and inducing R-loop accumulation. Nucleic Acids Res 47, 4068–4085. 10.1093/nar/gkz076.

39. Sessa, G., Gomez-Gonzalez, B., Silva, S., Perez-Calero, C., Beaurepere, R., Barroso, S., Martineau, S., Martin, C., Ehlen, A., Martinez, J.S., et al. (2021). BRCA2 promotes DNA-RNA hybrid resolution by DDX5 helicase at DNA breaks to facilitate their repairdouble dagger. EMBO J 40, e106018. 10.15252/embj.2020106018.

40. Yu, Z., Mersaoui, S.Y., Guitton-Sert, L., Coulombe, Y., Song, J., Masson, J.Y., and Richard, S. (2020). DDX5 resolves R-loops at DNA double-strand breaks to promote DNA repair and avoid chromosomal deletions. NAR Cancer 2, zcaa028. 10.1093/narcan/zcaa028.

41. Brickner, J.R., Garzon, J.L., and Cimprich, K.A. (2022). Walking a tightrope: The complex balancing act of R-loops in genome stability. Mol Cell 82, 2267–2297. 10.1016/j.molcel.2022.04.014.

42. Gatti, V., De Domenico, S., Melino, G., and Peschiaroli, A. (2023). Senataxin and R-loops homeostasis: multifaced implications in carcinogenesis. Cell Death Discov 9, 145. 10.1038/s41420-023-01441-x.

43. Cohen, S., Puget, N., Lin, Y.L., Clouaire, T., Aguirrebengoa, M., Rocher, V., Pasero, P., Canitrot, Y., and Legube, G. (2018). Senataxin resolves RNA:DNA hybrids forming at DNA double-strand breaks to prevent translocations. Nature communications 9, 533. 10.1038/s41467-018-02894-w.

44. He, W., and Parker, R. (2000). Functions of Lsm proteins in mRNA degradation and splicing. Curr Opin Cell Biol 12, 346–350. 10.1016/s0955-0674(00)00098-3.

45. Catala, R., Carrasco-Lopez, C., Perea-Resa, C., Hernandez-Verdeja, T., and Salinas, J. (2019). Emerging Roles of LSM Complexes in Posttranscriptional Regulation of Plant Response to Abiotic Stress. Front Plant Sci 10, 167. 10.3389/fpls.2019.00167.

46. Montemayor, E.J., Virta, J.M., Hayes, S.M., Nomura, Y., Brow, D.A., and Butcher, S.E. (2020). Molecular basis for the distinct cellular functions of the Lsm1-7 and Lsm2-8 complexes. Rna 26, 1400-1413. 10.1261/rna.075879.120.

47. Ingelfinger, D., Arndt-Jovin, D.J., Luhrmann, R., and Achsel, T. (2002). The human LSm1-7 proteins colocalize with the mRNA-degrading enzymes Dcp1/2 and Xrnl in distinct cytoplasmic foci. Rna 8, 1489–1501.

48. Reijns, M.A., Auchynnikava, T., and Beggs, J.D. (2009). Analysis of Lsm1p and Lsm8p domains in the cellular localization of Lsm complexes in budding yeast. The FEBS journal 276, 3602-3617. 10.1111/j.1742-4658.2009.07080.x.

49. Chowdhury, A., and Tharun, S. (2009). Activation of decapping involves binding of the mRNA and facilitation of the post-binding steps by the Lsm1-7-Pat1 complex. Rna 15, 1837–1848. 10.1261/rna.1650109.

50. Tharun, S. (2009). Lsm1-7-Pat1 complex: a link between 3’ and 5’-ends in mRNA decay? RNA Biol 6, 228-232. 10.4161/rna.6.3.8282.

51. Jones, C.I., Zabolotskaya, M.V., and Newbury, S.F. (2012). The 5’ --> 3’ exoribonuclease XRN1/Pacman and its functions in cellular processes and development. Wiley interdisciplinary reviews. RNA 3, 455–468. 10.1002/wrna.1109.

52. Mura, C., Cascio, D., Sawaya, M.R., and Eisenberg, D.S. (2001). The crystal structure of a heptameric archaeal Sm protein: Implications for the eukaryotic snRNP core. Proc Natl Acad Sci U S A 98, 5532–5537. 10.1073/pnas.091102298.

53. Liu, Y., Nomura, Y., Butcher, S.E., and Hoskins, A.A. (2025). RNA modifications and Prp24 coordinate Lsm2-8 binding dynamics during S. cerevisiae U6 snRNP assembly. J Biol Chem 301, 108497. 10.1016/j.jbc.2025.108497.

54. Arribas-Layton, M., Dennis, J., Bennett, E.J., Damgaard, C.K., and Lykke-Andersen, J. (2016). The C-Terminal RGG Domain of Human Lsm4 Promotes Processing Body Formation Stimulated by Arginine Dimethylation. Mol Cell Biol 36, 2226-2235. 10.1128/MCB.01102-15.

55. Gupta, R., Somyajit, K., Narita, T., Maskey, E., Stanlie, A., Kremer, M., Typas, D., Lammers, M., Mailand, N., Nussenzweig, A., et al. (2018). DNA Repair Network Analysis Reveals Shieldin as a Key Regulator of NHEJ and PARP Inhibitor Sensitivity. Cell 173, 972–988 e923. 10.1016/j.cell.2018.03.050.

56. Lafontaine, D.L.J., Riback, J.A., Bascetin, R., and Brangwynne, C.P. (2021). The nucleolus as a multiphase liquid condensate. Nat Rev Mol Cell Biol 22, 165–182. 10.1038/s41580-020-0272-6.

57. Li, H., Ju, Y., Liu, W.W., Ma, Y.Y., Ye, H., and Li, N. (2023). [Phase Separation of Purified Human LSM4 Protein]. Mol Biol (Mosk) 57, 124–126. 10.31857/S0026898423010068.

58. Shin, Y., Berry, J., Pannucci, N., Haataja, M.P., Toettcher, J.E., and Brangwynne, C.P. (2017). Spatiotemporal Control of Intracellular Phase Transitions Using Light-Activated optoDroplets. Cell 16*8*, 159-171 e114. 10.1016/j.cell.2016.11.054.

59. Singatulina, A.S., Hamon, L., Sukhanova, M.V., Desforges, B., Joshi, V., Bouhss, A., Lavrik, O.I., and Pastre, D. (2019). PARP-1 Activation Directs FUS to DNA Damage Sites to Form PARG-Reversible Compartments Enriched in Damaged DNA. Cell Rep 27, 1809–1821 e1805. 10.1016/j.celrep.2019.04.031.

60. Lombardi, S., Zilocchi, M., Nicsanu, R., and Barabino, S.M.L. (2025). Emerging connections: Poly(ADP-ribose), FET proteins and RNA in the regulation of DNA damage condensates. DNA Repair (Amst) 150, 103846. 10.1016/j.dnarep.2025.103846.

61. Chin Sang, C., Moore, G., Tereshchenko, M., Zhang, H., Nosella, M.L., Dasovich, M., Alderson, T.R., Leung, A.K.L., Finkelstein, I.J., Forman-Kay, J.D., and Lee, H.O. (2024). PARP1 condensates differentially partition DNA repair proteins and enhance DNA ligation. EMBO Rep 25, 5635–5666. 10.1038/s44319-024-00285-5.

62. Levone, B.R., Lenzken, S.C., Antonaci, M., Maiser, A., Rapp, A., Conte, F., Reber, S., Mechtersheimer, J., Ronchi, A.E., Muhlemann, O., et al. (2021). FUS-dependent liquid-liquid phase separation is important for DNA repair initiation. J Cell Biol 220. 10.1083/jcb.202008030.

63. Rhine, K., Dasovich, M., Yoniles, J., Badiee, M., Skanchy, S., Ganser, L.R., Ge, Y., Fare, C.M., Shorter, J., Leung, A.K.L., and Myong, S. (2022). Poly(ADP-ribose) drives condensation of FUS via a transient interaction. Mol Cell 82, 969–985 e911. 10.1016/j.molcel.2022.01.018.

64. Chappidi, N., Quail, T., Doll, S., Vogel, L.T., Aleksandrov, R., Felekyan, S., Kuhnemuth, R., Stoynov, S., Seidel, C.A.M., Brugues, J., et al. (2024). PARP1- DNA co-condensation drives DNA repair site assembly to prevent disjunction of broken DNA ends. Cell 187, 945–961 e918. 10.1016/j.cell.2024.01.015.

65. Teixeira, D., and Parker, R. (2007). Analysis of P-body assembly in Saccharomyces cerevisiae. Molecular biology of the cell 18, 2274–2287. 10.1091/mbc.e07-03-0199.

66. Chen, C.Y., Ezzeddine, N., and Shyu, A.B. (2008). Messenger RNA half-life measurements in mammalian cells. Methods in enzymology 448, 335–357. 10.1016/S0076-6879(08)02617-7.

67. Watson, M.J., and Thoreen, C.C. (2022). Measuring mRNA Decay with Roadblock-qPCR. Curr Protoc 2, e344. 10.1002/cpz1.344.

68. Watson, M.J., Park, Y., and Thoreen, C.C. (2021). Roadblock-qPCR: a simple and inexpensive strategy for targeted measurements of mRNA stability. Rna 27, 335–342. 10.1261/rna.076885.120.

69. Crossley, M.P., Brickner, J.R., Song, C., Zar, S.M.T., Maw, S.S., Chedin, F., Tsai, M.S., and Cimprich, K.A. (2021). Catalytically inactive, purified RNase H1: A specific and sensitive probe for RNA-DNA hybrid imaging. J Cell Biol 220. 10.1083/jcb.202101092.

70. Pinder, J., Salsman, J., and Dellaire, G. (2015). Nuclear domain ’knock-in’ screen for the evaluation and identification of small molecule enhancers of CRISPR-based genome editing. Nucleic Acids Res 43, 9379–9392. 10.1093/nar/gkv993.

71. Mattout, A., Gaidatzis, D., Padeken, J., Schmid, C.D., Aeschimann, F., Kalck, V., and Gasser, S.M. (2020). LSM2-8 and XRN-2 contribute to the silencing of H3K27me3-marked genes through targeted RNA decay. Nat Cell Biol 22, 579–590. 10.1038/s41556-020-0504-1.

72. Perea-Resa, C., Carrasco-Lopez, C., Catala, R., Tureckova, V., Novak, O., Zhang, W., Sieburth, L., Jimenez-Gomez, J.M., and Salinas, J. (2016). The LSM1-7 Complex Differentially Regulates Arabidopsis Tolerance to Abiotic Stress Conditions by Promoting Selective mRNA Decapping. Plant Cell 28, 505–520. 10.1105/tpc.15.00867.

73. Perea-Resa, C., Hernandez-Verdeja, T., Lopez-Cobollo, R., del Mar Castellano, M., and Salinas, J. (2012). LSM proteins provide accurate splicing and decay of selected transcripts to ensure normal Arabidopsis development. Plant Cell *24*, 4930-4947. 10.1105/tpc.112.103697.

74. Li, P., Qi, F., Zhu, W., Li, J., Shi, J., Tu, X., Wang, M., Chen, P., Liu, B.F., and Li, Y. (2026). High-throughput identification of endogenous biomolecular condensates and phase-separating proteins. Nat Protoc. 10.1038/s41596-025-01327-5.

75. Murthy, A.C., Dignon, G.L., Kan, Y., Zerze, G.H., Parekh, S.H., Mittal, J., and Fawzi, N.L. (2019). Molecular interactions underlying liquid-liquid phase separation of the FUS low-complexity domain. Nat Struct Mol Biol 26, 637-648. 10.1038/s41594-019-0250-x.

76. Tangudu, N.K., and Aird, K.M. (2022). 53BP1: guarding the genome with a novel liquid weapon. Commun Biol 5, 435. 10.1038/s42003-022-03401-0.

77. Zhang, L., Geng, X., Wang, F., Tang, J., Ichida, Y., Sharma, A., Jin, S., Chen, M., Tang, M., Pozo, F.M., et al. (2022). 53BP1 regulates heterochromatin through liquid phase separation. Nature communications 13, 360. 10.1038/s41467-022-28019-y.

78. Morales, J.C., Richard, P., Patidar, P.L., Motea, E.A., Dang, T.T., Manley, J.L., and Boothman, D.A. (2016). XRN2 Links Transcription Termination to DNA Damage and Replication Stress. PLoS Genet 12, e1006107. 10.1371/journal.pgen.1006107.

79. Villarreal, O.D., Mersaoui, S.Y., Yu, Z., Masson, J.Y., and Richard, S. (2020). Genome-wide R-loop analysis defines unique roles for DDX5, XRN2, and PRMT5 in DNA/RNA hybrid resolution. Life Sci Alliance *3*. 10.26508/lsa.202000762.

80. Krishnan, R., Lapierre, M., Gautreau, B., Nixon, K.C.J., El Ghamrasni, S., Patel, P.S., Hao, J., Yerlici, V.T., Guturi, K.K.N., St-Germain, J., et al. (2023). RNF8 ubiquitylation of XRN2 facilitates R-loop resolution and restrains genomic instability in BRCA1 mutant cells. Nucleic Acids Res 51, 10484–10505. 10.1093/nar/gkad733.

81. Brannan, K., Kim, H., Erickson, B., Glover-Cutter, K., Kim, S., Fong, N., Kiemele, L., Hansen, K., Davis, R., Lykke-Andersen, J., and Bentley, D.L. (2012). mRNA decapping factors and the exonuclease Xrn2 function in widespread premature termination of RNA polymerase II transcription. Mol Cell 46, 311-324. 10.1016/j.molcel.2012.03.006.

82. Mugridge, J.S., and Gross, J.D. (2013). Judge, jury, and executioner: DXO functions as a decapping enzyme and exoribonuclease in pre-mRNA quality control. Mol Cell 50, 2–4. 10.1016/j.molcel.2013.03.025.

83. Domingo-Prim, J., Endara-Coll, M., Bonath, F., Jimeno, S., Prados-Carvajal, R., Friedlander, M.R., Huertas, P., and Visa, N. (2019). EXOSC10 is required for RPA assembly and controlled DNA end resection at DNA double-strand breaks. Nature communications 10, 2135. 10.1038/s41467-019-10153-9.

84. He, G., Gu, K., Wei, J., and Zhang, J. (2024). METTL3-mediated the m6A modification of SF3B4 facilitates the development of non-small cell lung cancer by enhancing LSM4 expression. Thorac Cancer 15, 919–928. 10.1111/1759-7714.15275.

85. Sun, X., Zhang, J., Xiao, C., and Ge, Z. (2022). Expression profile and prognostic values of LSM family in skin cutaneous melanoma. BMC Med Genomics 15, 238. 10.1186/s12920-022-01395-6.

86. Ren, Z., Li, Y., Yang, X., Zhang, N., Wang, F., and He, Z. (2025). LSM4 as a potential prognostic indicator and therapeutic target in triple-negative breast cancer progression. Gland Surg 14, 1990–2004. 10.21037/gs-2025-48.

87. Sun, Z.P., Tan, Z.G., and Peng, C. (2022). Long noncoding RNA LINC01419 promotes hepatocellular carcinoma malignancy by mediating miR-485-5p/LSM4 axis. Kaohsiung J Med Sci 38, 826–838. 10.1002/kjm2.12566.

88. Chen, L., Lin, Y.H., Liu, G.Q., Huang, J.E., Wei, W., Yang, Z.H., Hu, Y.M., Xie, J.H., and Yu, H.Z. (2021). Clinical Significance and Potential Role of LSM4 Overexpression in Hepatocellular Carcinoma: An Integrated Analysis Based on Multiple Databases. Front Genet 12, 804916. 10.3389/fgene.2021.804916.

89. Ta, H.D.K., Wang, W.J., Phan, N.N., An Ton, N.T., Anuraga, G., Ku, S.C., Wu, Y.F., Wang, C.Y., and Lee, K.H. (2021). Potential Therapeutic and Prognostic Values of LSM Family Genes in Breast Cancer. Cancers (Basel) 13. 10.3390/cancers13194902.

90. Yin, J., Lin, C., Jiang, M., Tang, X., Xie, D., Chen, J., and Ke, R. (2021). CENPL, ISG20L2, LSM4, MRPL3 are four novel hub genes and may serve as diagnostic and prognostic markers in breast cancer. Sci Rep *11*, 15610. 10.1038/s41598-021-95068-6.

91. Hou, W., and Zhang, Y. (2021). Circ_0025033 promotes the progression of ovarian cancer by activating the expression of LSM4 via targeting miR-184. Pathol Res Pract 217, 153275. 10.1016/j.prp.2020.153275.

92. Bishara, L.A., Abu-Zhayia, E.R., Nicola, M., and Ayoub, N. (2025). C8orf33 dictates DNA double-strand break repair choice by modulating KAT8-mediated H4K16 acetylation. Cell Death Dis 16, 834. 10.1038/s41419-025-08194-8.

93. Khoury-Haddad, H., Guttmann-Raviv, N., Ipenberg, I., Huggins, D., Jeyasekharan, A.D., and Ayoub, N. (2014). PARP1-dependent recruitment of KDM4D histone demethylase to DNA damage sites promotes double-strand break repair. Proc Natl Acad Sci U S A 111, E728–737. 10.1073/pnas.1317585111.

94. Kim, D., Langmead, B., and Salzberg, S.L. (2015). HISAT: a fast spliced aligner with low memory requirements. Nat Methods 12, 357–360. 10.1038/nmeth.3317.

95. Love, M.I., Huber, W., and Anders, S. (2014). Moderated estimation of fold change and dispersion for RNA-seq data with DESeq2. Genome biology 15, 550. 10.1186/s13059-014-0550-8.

96. Ge, S.X., Jung, D., and Yao, R. (2020). ShinyGO: a graphical gene-set enrichment tool for animals and plants. Bioinformatics 36, 2628-2629. 10.1093/bioinformatics/btz931.

97. Shen, S., Park, J.W., Lu, Z.X., Lin, L., Henry, M.D., Wu, Y.N., Zhou, Q., and Xing, Y. (2014). rMATS: robust and flexible detection of differential alternative splicing from replicate RNA-Seq data. Proc Natl Acad Sci U S A 111, E5593–5601. 10.1073/pnas.1419161111.

98. Katoh, K., Rozewicki, J., and Yamada, K.D. (2019). MAFFT online service: multiple sequence alignment, interactive sequence choice and visualization. Brief Bioinform 20, 1160–1166. 10.1093/bib/bbx108.

99. Katoh, K., and Standley, D.M. (2013). MAFFT multiple sequence alignment software version 7: improvements in performance and usability. Mol Biol Evol 30, 772–780. 10.1093/molbev/mst010.

100. Culjkovic-Kraljacic, B., Skrabanek, L., Revuelta, M.V., Gasiorek, J., Cowling, V.H., Cerchietti, L., and Borden, K.L.B. (2020). The eukaryotic translation initiation factor eIF4E elevates steady-state m(7)G capping of coding and noncoding transcripts. Proc Natl Acad Sci U S A 117, 26773–26783. 10.1073/pnas.2002360117.

101. Langmead, B., and Salzberg, S.L. (2012). Fast gapped-read alignment with Bowtie 2. Nat Methods 9, 357-359. 10.1038/nmeth.1923.

102. Li, H., Handsaker, B., Wysoker, A., Fennell, T., Ruan, J., Homer, N., Marth, G., Abecasis, G., Durbin, R., and Genome Project Data Processing, S. (2009). The Sequence Alignment/Map format and SAMtools. Bioinformatics *25*, 2078-2079. 10.1093/bioinformatics/btp352.

103. Ramirez, F., Ryan, D.P., Gruning, B., Bhardwaj, V., Kilpert, F., Richter, A.S., Heyne, S., Dundar, F., and Manke, T. (2016). deepTools2: a next generation web server for deep-sequencing data analysis. Nucleic Acids Res 44, W160–165. 10.1093/nar/gkw257.

