## Supplemental Information for "LSm4 biomolecular condensates drive XRN2-mediated RNA decay at DNA double-strand breaks to facilitate repair"

This PDF file includes supplementary Figures and Legends: S1 to S5

**Supplementary Figures:**

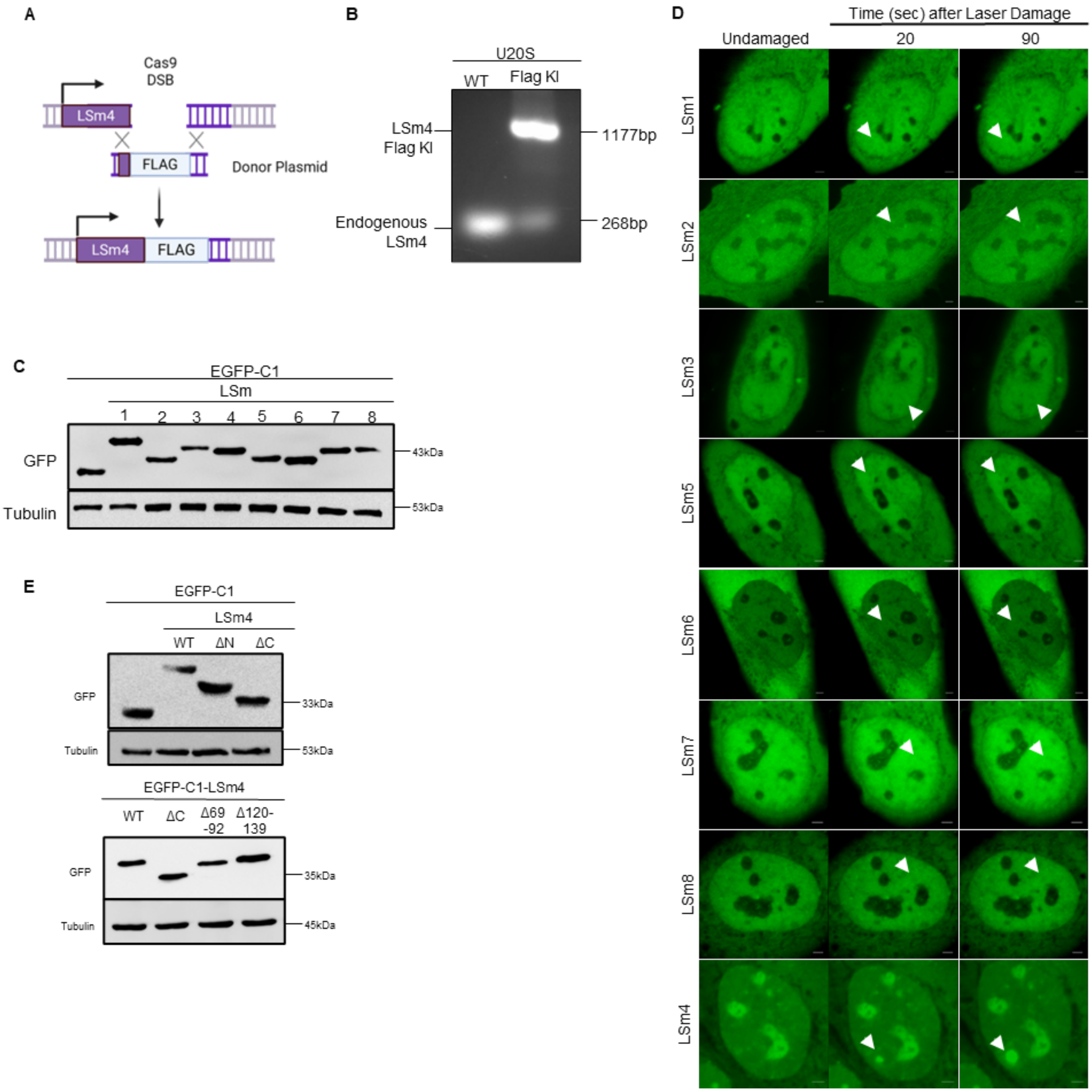


**Figure S1. Related to Figure 1: γH2AX enrichment at *Asi*SI-induced DSBs**
**(A)** A schematic describing DSB induction at *Asi*SI sites following 4OHT treatment in U2OS-DIvA cells. **(B)** Average profile of γH2AX enrichment between 4OHT-treated and untreated DIvA cells in 6Mb window surrounding 214 *Asi*SI sites with highest γH2AX (left) or 214 random genomic loci (right). Top: values are expressed as average normalized ChIP-seq reads. Bottom: values are expressed as average log2 ratio between 4OHT-treated and untreated cells.

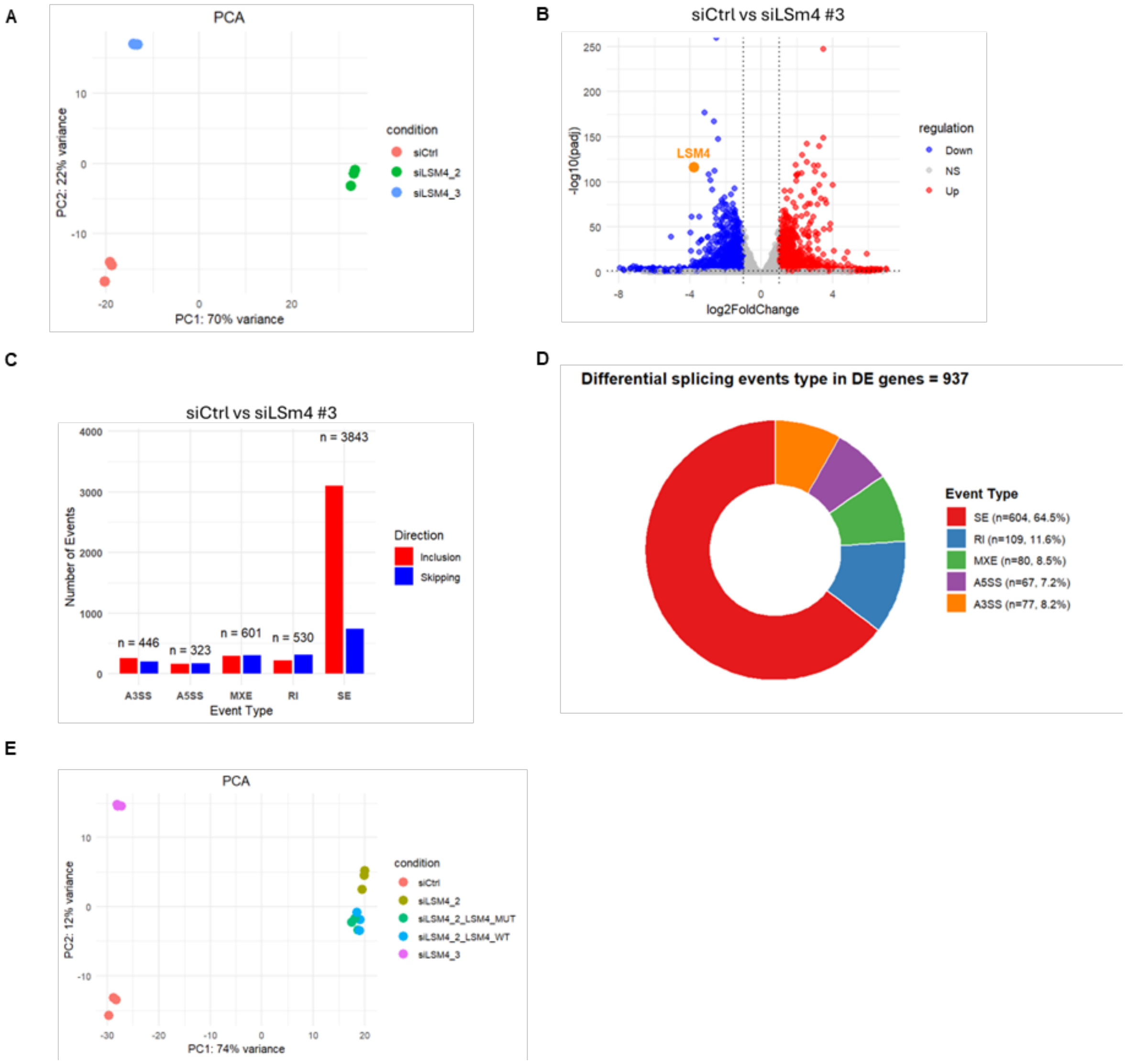


**Figure S2. Related to Figure 4. LSm4 regulates RNA decapping and decay independently of its roles in transcription and splicing.** (A) Principal component analysis (PCA) of RNA-seq data shows a high degree of reproducibility among the replicate samples within each group. (B) Volcano plot summarizing differential gene expression data obtained from RNA-seq analysis between control and siLSm4 #3-transfected U2OS cells. Upregulated genes with log2FoldChange(siLSm4 #3/control) > 1 and *P*adj-value < 0.05 are marked in red, while downregulated genes with log2FoldChange(siLSm4 #3/control) < -1 and *P*adj-value < 0.05 are marked in blue. (C) Summary of significant alternative splicing events leading to skipping (blue) or inclusion (red) observed upon siLSm4 #3 treatment as detected by rMATS. Significantly altered splicing events were classified as having a minimum inclusion level difference of 0.1, *p* value < 0.01, and FDR < 0.01. SE: skipped exon, MXE: mutually exclusive exons, A5SS: alternative 5’ splice site, A3SS: alternative 3’ splice site, IR: intron retention. (D) Donut chart showing the distribution of significant alternative splicing events among the spliced DE genes (identified in Figure 4E). (A) PCA of RNA-seq data including siLSm4 #2 + LSm4$^{WT}$ and siLSm4 #2 + LSm4$^{\Delta 69-92}$ samples.

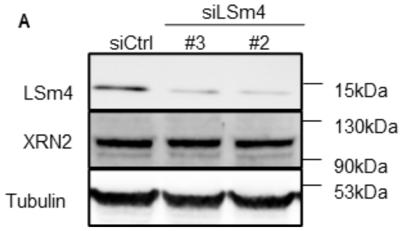


**Figure S3. Related to Figure 5: LSm4 depletion does not affect XRN2 levels.** (A) Immunoblot analysis of XRN2 levels in control and LSm4-depleted U2OS-DIvA cells.

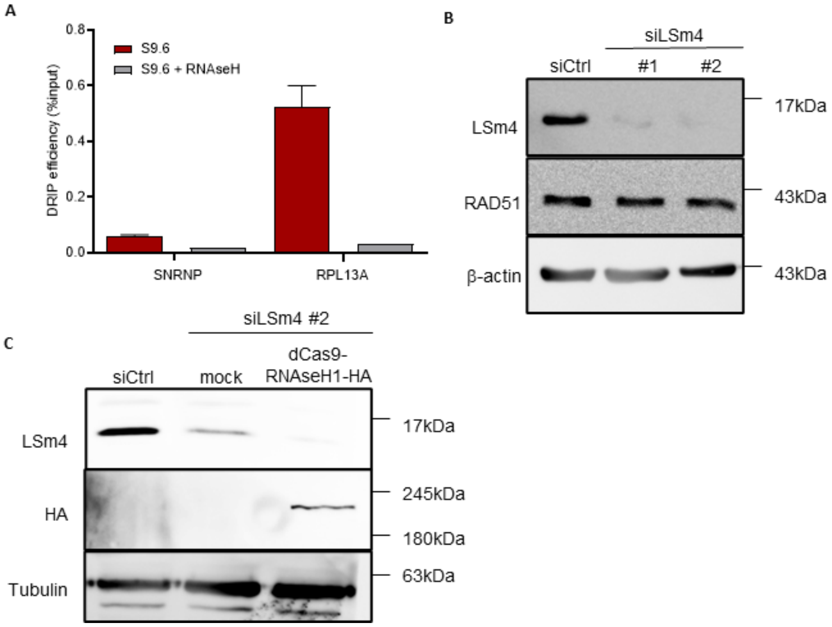

**Figure S4. Related to Figure 6: Validations of DRIP efficiency and LSm4 expression in cells.** (A) DRIP-qPCR for two genomic loci devoid (*SNRNP*) or enriched (*RPL13A*) in R-loops, performed in U2OS-DIvA cells in the presence or absence of RNaseH1 treatment. (B) Immunoblot analysis for RAD51 levels in LSm4-proficient and deficient U2OS-DIvA cells. (C) Immunoblot analysis for LSm4 and HA in LSm4-proficient and deficient U2OS-DIvA cells. HA represents the expression of dCas9-RNaseH1-HA.

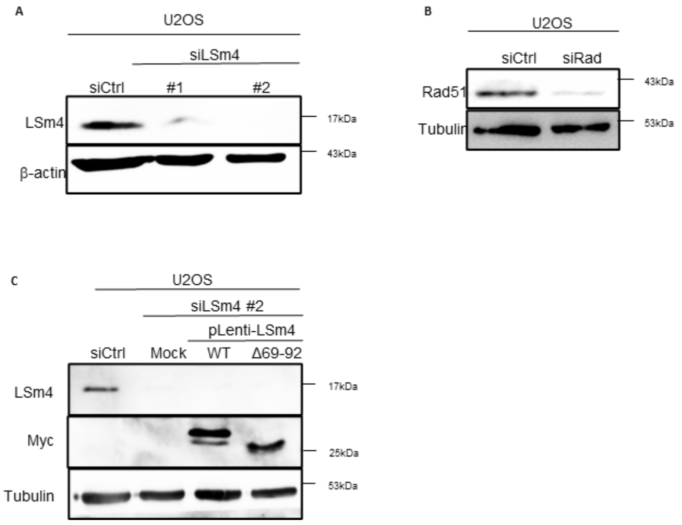


**Figure S5. Related to Figure 7: Validation of LSm4 and RAD51 levels.** (A) Immunoblot analysis for LSm4 levels in U2OS cells. (B) Immunoblot analysis for RAD51 levels in U2OS cells. (C) Immunoblot analysis for LSm4 and Myc levels in U2OS cells expressing either siCtrl or siLSm4 #2, stably expressing myc-LSm4$^{WT}$ or myc-LSm4$^{\Delta 69–92}$.

**Supplementary Movie 1:** Time-lapse movie showing Optodroplet formation of Cry2WT following exposure to Blue Light, Related to Figure 1D.

**Supplementary Movie 2:** Time-lapse movie showing Optodroplet formation of LSm4-Cry2WT following exposure to Blue Light, Related to Figure 1D.

**Supplementary Movie 3:** Time-lapse movie showing Optodroplet formation of LSm1-Cry2WT following exposure to Blue Light, Related to Figure 1D.

**Supplementary Movie 4:** Time-lapse movie showing Optodroplet formation of LSm8-Cry2WT following exposure to Blue Light, Related to Figure 1D.

**Supplementary Movie 5:** Time-lapse movie showing Optodroplet formation of LSm4$^{\Delta 69–92}$-Cry2WT following exposure to Blue Light, Related to Figure 1H.
